# Loss of NR2E3 disrupts rod photoreceptor cell maturation causing a fate switch late in human retinal development

**DOI:** 10.1101/2023.06.30.547279

**Authors:** Nathaniel K. Mullin, Laura R. Bohrer, Andrew P. Voigt, Lola P. Lozano, Allison Wright, Robert F. Mullins, Edwin M. Stone, Budd A. Tucker

## Abstract

While dysfunction and death of light-detecting photoreceptor cells underlie most inherited retinal dystrophies, knowledge of the species-specific details of human rod and cone photoreceptor cell development remains limited. Here, we generate retinal organoids using induced pluripotent stem cells (iPSC) derived from a patient with genetic photoreceptor disease due to mutations in *NR2E3*, an isogenic control, and an unrelated control. Organoids were sampled using single-cell RNA sequencing across the developmental window encompassing photoreceptor specification, emergence, and maturation, up to 260 days of *in vitro* differentiation. Using single-cell transcriptomics data, we reconstruct the rod photoreceptor developmental lineage and identify a branchpoint in development unique to the disease state that gives rise to a divergent rod photoreceptor cell population. We show that the rod-specific transcription factor NR2E3 is required for the proper expression of genes involved in phototransduction, including expression of the light-sensitive protein rhodopsin, which is absent in divergent rods. NR2E3-null rods additionally misexpress several cone-specific phototransduction genes at both the transcript and protein level. Using joint multimodal single-cell sequencing on late-stage retinal organoids, we further identify specific putative regulatory sites where rod-specific factors act to steer rod and cone photoreceptor cell development. Importantly, these findings are strikingly different than those observed in rodent models of disease. Together, these data provide a roadmap of human photoreceptor development and leverage patient iPSCs to define the specific roles of rod transcription factors in photoreceptor cell emergence and maturation.

## INTRODUCTION

The human retina is a transparent multilayered neural tissue lining the posterior two-thirds of the eye that is responsible for detecting and sending visual information to the brain via the second cranial nerve. Light detecting photoreceptor cells, which make up the outermost layer of the neural retina, emerge during early development from a pool of multipotent neural progenitors (Morrow et al., 1998; Trimarchi et al., 2008). In humans, death of photoreceptor cells, which is associated with inherited retinal degenerative disease, is a major cause of incurable blindness. Disease causing variants in genes such as *USH2A*, *RHO*, and *RPGR*, which are the leading causes of recessive, dominant, and x-linked disease respectively, result in progressive vision loss typically beginning in late teens and young adults (Stone et al., 2017). The relatively late onset and slowly progressive nature of these disorders allow for real time investigation using a variety of different clinical approaches. For transcription factor genes that regulate photoreceptor cell fate commitment and maturation, clinical evaluation of the early disease state is often impossible. That is, loss of function mutations in transcription factor genes often result in the absence of specific cell types at birth (Cvekl and Callaerts, 2017; Diacou et al., 2022; Hull et al., 2014; Lima Cunha et al., 2019; Sun and Chen, 2023). Understanding the precise role of transcription factors in human photoreceptor cell development and how loss of function mutations cause disease has the possibility to suggest novel approaches to rescue dysfunctional photoreceptors in patients diagnosed with an inherited retinal degeneration.

Mutations in the nuclear receptor subfamily 2 group E member 3 (*NR2E3*) gene cause Enhanced S Cone Syndrome (ESCS) (OMIM 268100), a congenital retinal disease characterized by night blindness, hypersensitivity to short-wavelength light, and eventual loss of visual acuity (Haider et al., 2000; Hood et al., 1995). Rod photoreceptors are the primary effectors of vision in dim light. NR2E3 is expressed in rod photoreceptors and plays an essential role in rod development in concert with upstream rod transcription factors including Neural Retina Leucine Zipper (NRL) (Mears et al., 2001). The retinas of ESCS patients additionally exhibit disorganization of the normal cellular layers (Milam et al., 2002). Postmortem histological observation of ESCS patients’ retinas have demonstrated a lack of staining for rhodopsin, the primary functional molecule of light sensitivity in rod photoreceptors (Milam et al., 2002). Additionally, there are an increased proportion of cones, specifically S-cones, which mislocalize into the layers of the retina typically restricted to rod photoreceptors (Milam *et al*., 2002). Much of the knowledge of mammalian rod photoreceptor specification comes from murine models (Brzezinski and Reh, 2015). The Rd7 mouse harbors spontaneous mutations in *Nr2e3*, which cause a retinal degeneration phenotype reminiscent of the ESCS patients (Iannaccone et al., 2021). However, key rod function genes such as the light-sensitive protein rhodopsin (*Rho*) are expressed in the Nr2e3-deficient mouse retina (Aísa-Marín et al., 2023; Corbo and Cepko, 2005; Haider et al., 2006; Iannaccone *et al*., 2021), in contrast to the complete loss of rod function observed in ESCS patients. As the mouse has a rod-dominant retina lacking a cone-rich macula (Volland et al., 2015), and the requirement of core rod transcription factors for rod specification is known to vary in other vertebrates (Oel et al., 2020), the precise regulatory processes governing rod and cone photoreceptor specification and maturation may differ between species. As such, it is essential to study the specific clinically relevant targets in human retinal cells to understand the pathogenesis of *NR2E3* associated disease and related conditions.

To define the specific timing and targets of NR2E3 activity in human retinal development, we performed transcriptome, chromatin, and protein-level analysis across a 260-day time course from early retinal commitment through photoreceptor cell maturation. We utilized an effectively NR2E3-null human iPSC line derived from a clinically diagnosed ESCS patient. To control for the effects of genetic background on organoid differentiation efficiency, we differentiated two control lines in parallel: a CRISPR-corrected isogenic control and an unrelated healthy donor control. We used single-cell RNA sequencing to identify a population of rod photoreceptor cells (which we categorize as “divergent”) that emerged between differentiation day 80 and 120 in the NR2E3-null organoids. We showed that these cells, which persist throughout retinal development, co-express rod and cone photoreceptor cell markers that are typically expressed in mature cells at both the transcript and protein level. Using single-cell multimodal sequencing, we showed that misregulation of these genes is due to changes in the activity of cis-regulatory elements following loss of NR2E3 function. Interestingly, at later stages of retinal development an increase in the number of blue cones at the expense of divergent rods was detected. That is, while divergent rods persisted throughout development, their proportion decreased as the proportion of blue cones increased, which may suggest divergent rod to blue cone conversion. Together, these data define a specific role for *NR2E3* in human photoreceptor cell development that appears to be distinct from that of rodents, which sheds light on the cellular changes underlying the ESCS retinal phenotype.

## RESULTS

### Retinal organoids produce developmentally timed cell types

To determine how and when NR2E3 acts in human retinal development, we used a previously described iPSC line (Bohrer et al., 2019) derived from an ESCS patient with a homozygous c.119-2A>C mutation in *NR2E3*. This mutation causes the inclusion of a portion of intron 1 which creates a frame shift and premature stop codon following the first exon, rendering it null (referred to as NR2E3-null going forward). We showed that monoallelic correction of c. 119-2A>C in patient-derived iPSCs by CRISPR-mediated homology dependent repair (HDR) restores the ability of photoreceptor cells to make wild-type *NR2E3* transcript during retinal cell differentiation (Bohrer, 2019). To capture both developing and terminally differentiated cell types, retinal organoids were generated from no disease control (ND control), NR2E3-null, and CRISPR-corrected NR2E3 (isogenic control) and these organoids were initially sampled across a 160-day time course (**Figure 1A**). Organoids were assayed using single-cell transcriptome profiling and immunohistochemistry (**Figure 1, Figure S1, Figure S2**). Data from cells collected on days 40, 80, 120, and 160 (hereafter referred to as D40, D80, D120, and D160) of differentiation across all three lines were aggregated and annotated using previously published human organoid and fetal retina scRNAseq data (Sridhar et al., 2020). Cell type emergence followed the known developmental cadence of retinal formation (Morrow et al., 1998), with progenitors giving rise to cone photoreceptors and inner retinal cells first, followed by waves of rod photoreceptor and bipolar cell emergence (**Figure 1B**). Notably, all expected cell types, including rod photoreceptors, were observed in each line (**Figure 1C-E**).

**Figure 1.**
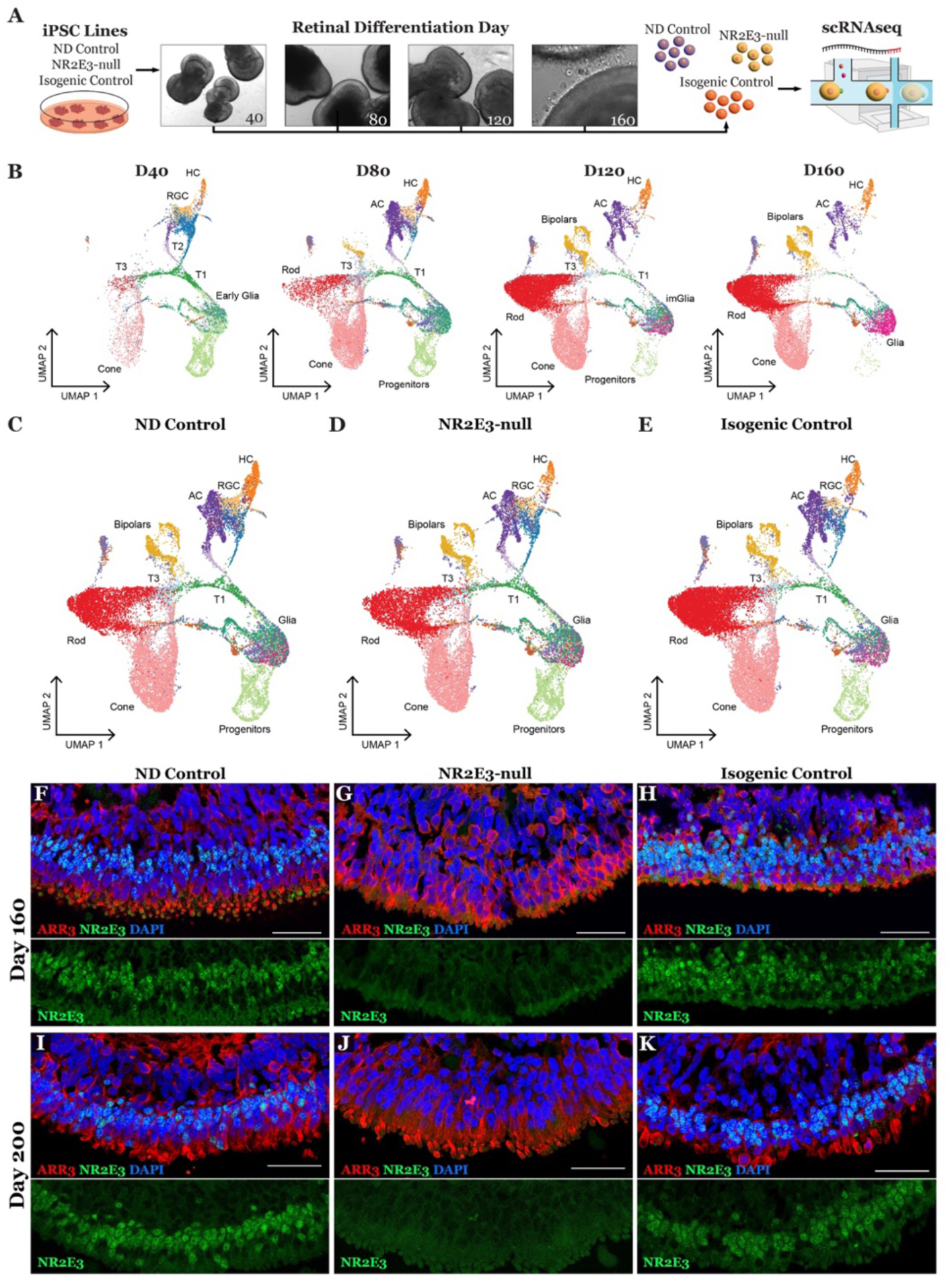
Modeling pathological retinal development using retinal organoids. **A)** Experimental schematic for organoid differentiation time course with scRNAseq**. B)** Integrated and annotated cells recovered from all scRNAseq samples projected in 2D space using UMAP embeddings. Cells are shown grouped by timepoint of collection, highlighting normal developmental timing**. C-E)** Cells from all timepoints are shown split by cell line of origin. All cell lines generate cells of each annotated type. **F)** ND control organoids express NR2E3 (green) in rod nuclei at D160 of differentiation. Cone photoreceptors express cone arrestin (ARR3, red). **G)** NR2E3-null organoids express cone arrestin but lack expression of NR2E3. This confirms that biallelic c.119-2A>C mutations are sufficient to cause loss of protein expression in the organoid model. **H)** Monoallelic correction of NR2E3 restores expression of NR2E3 in D160 organoids. **I-K)** At D200 of differentiation, NR2E3 expression remains high in ND control and isogenic control lines and is absent in NR2E3-null samples. Scalebar = 50um.

NR2E3 is required for normal rod photoreceptor development (Cheng et al., 2006; Cheng et al., 2004; Milam et al., 2002) and is known to be expressed soon after the induction of its upstream activator, NRL (Bumsted O’Brien et al., 2004). We observed the emergence of rod photoreceptors in organoids by D120 of differentiation in all three lines (**Figure 1B - E**). We next stained for the NR2E3 protein in fixed sections of retinal organoids at comparable timepoints. Control organoids expressed NR2E3 in the nucleus at D160 (**Figure 1F**). No NR2E3 protein was detected in the NR2E3-null organoids at the same timepoint (**Figure 1G**) and monoallelic correction of the locus restored normal expression (**Figure 1H**). The same pattern of protein expression persisted to D200 (**Figure 1I - K**), indicating that the c.119-2A>C *NR2E3* mutations cause a total lack, rather than delay, of NR2E3 protein expression in human retinal organoids.

### Divergent rods emerge in NR2E3-null organoids following rod commitment

Since NR2E3 is known to be required for rod photoreceptor cell formation, we next asked how NR2E3-null rods differed transcriptionally from normal rod photoreceptors. We computationally isolated the photoreceptor lineage within the dataset to enable comparison of developmental lineages of rod and cone photoreceptors. D40, D80, D120, and D160 cells from all three lines that were annotated as Progenitors, T1, T3, Cone, or Rod (i.e. from **Figure 1B**) were reprocessed using potential of heat diffusion for affinity-based transition embedding (PHATE) (Moon et al., 2019), a dimensionality-reduction technique suited to maintaining the branching structure in developmental datasets. The ordering of cells in the PHATE embedding matched collection timepoints of the samples (**Figure 2A**), lending confidence to the biological relevance of this approach. Cells were re-clustered and manually re-annotated based on the PHATE embedding (**Figure 2B**) using timepoint and expression information of marker genes (**Figure S3A**). In addition to refining the maturity of rod and cone photoreceptors (e.g., early cone, immature cone, cone), a novel cluster was also observed in the PHATE reduction that appeared largely restricted to the NR2E3-null cells (**Figure 2C**). Since it branched from the early rod cluster, we named these cells “divergent rods”.

**Figure 2.**
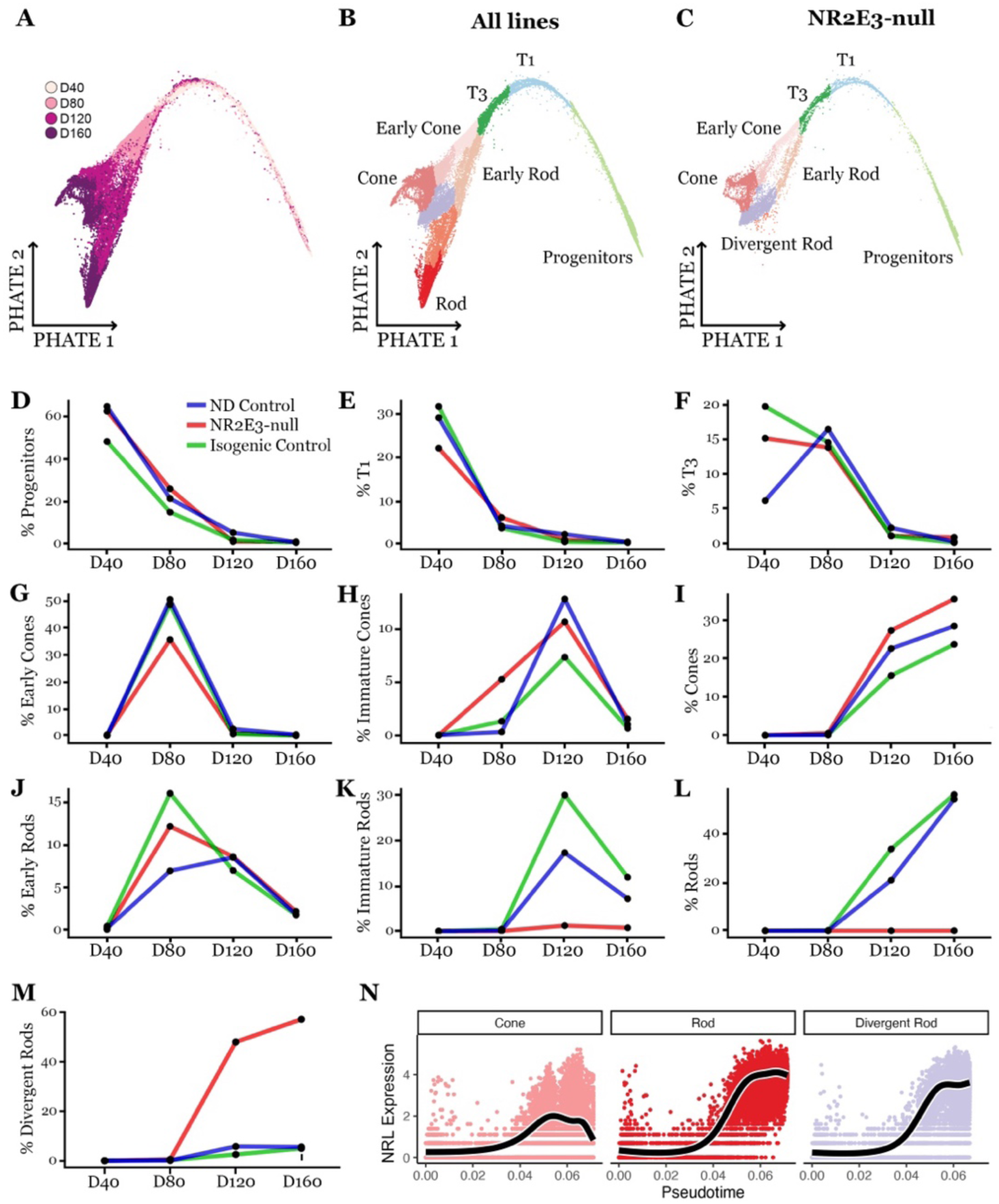
Divergent rods emerge in NR2E3-null organoids. **A)** Cells within the photoreceptor lineage projected in low dimensional space using PHATE embeddings. Cells are colored by timepoint of sample collection showing preservation of temporal order by PHATE reduction. **B)** Cells annotated based on timepoint and PHATE-derived cluster. Cells from both NR2E3-null and control lines are shown together. **C)** Cells annotated based on PHATE clustering from only the NR2E3-null line are shown. No immature rod or rod photoreceptors are present, and a large divergent rod population is present. **D – F)** The proportion of progenitors and intermediate progenitors (i.e., T1 and T3) decreases uniformly across differentiation of control and NR2E3-null lines. **G – I)** The proportion of maturing cones follows differentiation timepoint in all three lines with no major deviations. **J)** All three lines form early rod photoreceptors at D80. **K, L)** Only ND control and isogenic control lines form immature and mature rod photoreceptors at D120 and D160. M) Divergent rods emerge by D120 and are largely restricted to the NR2E3-null line. **N)** The PHATE embeddings from **B** were used to generate trajectories capturing cone, rod, and divergent rod development. *NRL* expression is plotted against pseudotime for each lineage on a log scale. *NRL* expression is observed at comparable levels in rod and divergent rod lineages and is induced at the same point in pseudotime.

To better understand the developmental origin of the divergent rod cluster, we calculated the proportion of each cell type at each timepoint of photoreceptor lineage differentiation (**Figure 2D - M**). When cell type proportions are plotted across time, the disappearance of progenitors followed by the emergence of maturing photoreceptor cells is seen (**Figure 2D-L**), indicating that proper commitment and maturation of cell types occur within retinal organoids of all three lines. However, emergence of normally mature rods is observed only in control lines (**Figure 2K, L**) while formation of divergent rods is restricted to the NR2E3-null line (red line) at D120 (**Figure 2M**). Notably, NR2E3-null cells produce early rods at D80 at roughly similar proportions to control lines (**Figure 2J**), counter to the hypothesis that NR2E3-null photoreceptor progenitors would be uniformly shunted into a cone cell fate at this juncture. Instead, these data support rod malformation in NR2E3-null occurring after rod photoreceptor cell fate commitment at D80.

To further investigate the differences between normal and pathological rod differentiation, we next identified three trajectories through the PHATE embedding using Slingshot (Street et al., 2018) (**Figure S3B-E**). These trajectories describe the maturation of progenitor cells into normal cones, normal rods, or divergent rods. Such trajectories can be used to compute pseudotime values for each cell within each lineage enabling downstream gene expression analysis and visualization (**Figure S3C-H**). We plotted expression of NRL against pseudotime within each lineage (**Figure 2N**) and observe comparable timing and level of NRL expression in the normal and divergent rod lineages. NRL levels are markedly lower in the cone lineage. This finding suggests that NRL-expressing divergent rods initially develop normally and that NR2E3 absence causes a failure in rod maturation later in development.

### Joint multimodal sequencing of divergent rod transcriptome and chromatin accessibility

To confirm that the emergence of divergent rods was not an artifact of batch-to-batch variability of organoid differentiation and to gain information on chromatin remodeling following NR2E3 loss in rods, we performed single-nucleus multimodal sequencing on retinal organoid nuclei isolated from an independent round of differentiation (**Figure 3A**) of the NR2E3-null and isogenic control lines. Nuclei were collected from timepoints after the emergence of divergent rods (D160 and D260). Joint multimodal RNA and ATAC sequencing were performed to query the differential gene expression and accessibility of regulatory regions in NR2E3-null rods. First, nuclei were clustered on a weighted combination of ATAC and gene expression modalities (**Figure 3B**). Clusters were annotated using gene expression data of canonical marker genes (**Figure 3B**). Cone and rod photoreceptor nuclei were captured in both NR2E3-null and isogenic control organoids at D160 and D260, recapitulating the finding of our previous time course study (**Figures 1 & 2**). Using the D40-D160 gene expression data from the photoreceptor lineage (**Figure 2B-C**), rod, cone, and divergent rod gene modules were computed (**Figure S4A-J**). Nuclei in the multimodal dataset were scored for each module. An enrichment within the NR2E3-null rod cluster for the divergent rod gene module was observed (**Figure 3C, Figure S4G-I**). This enrichment was not observed in isogenic control rods (**Figure 3C, Figure S4H, I**). These data show that the emergence of divergent rods in NR2E3-null organoids is reproducible and robust to discovery across different sequencing modalities and rounds of differentiation.

**Figure 3.**
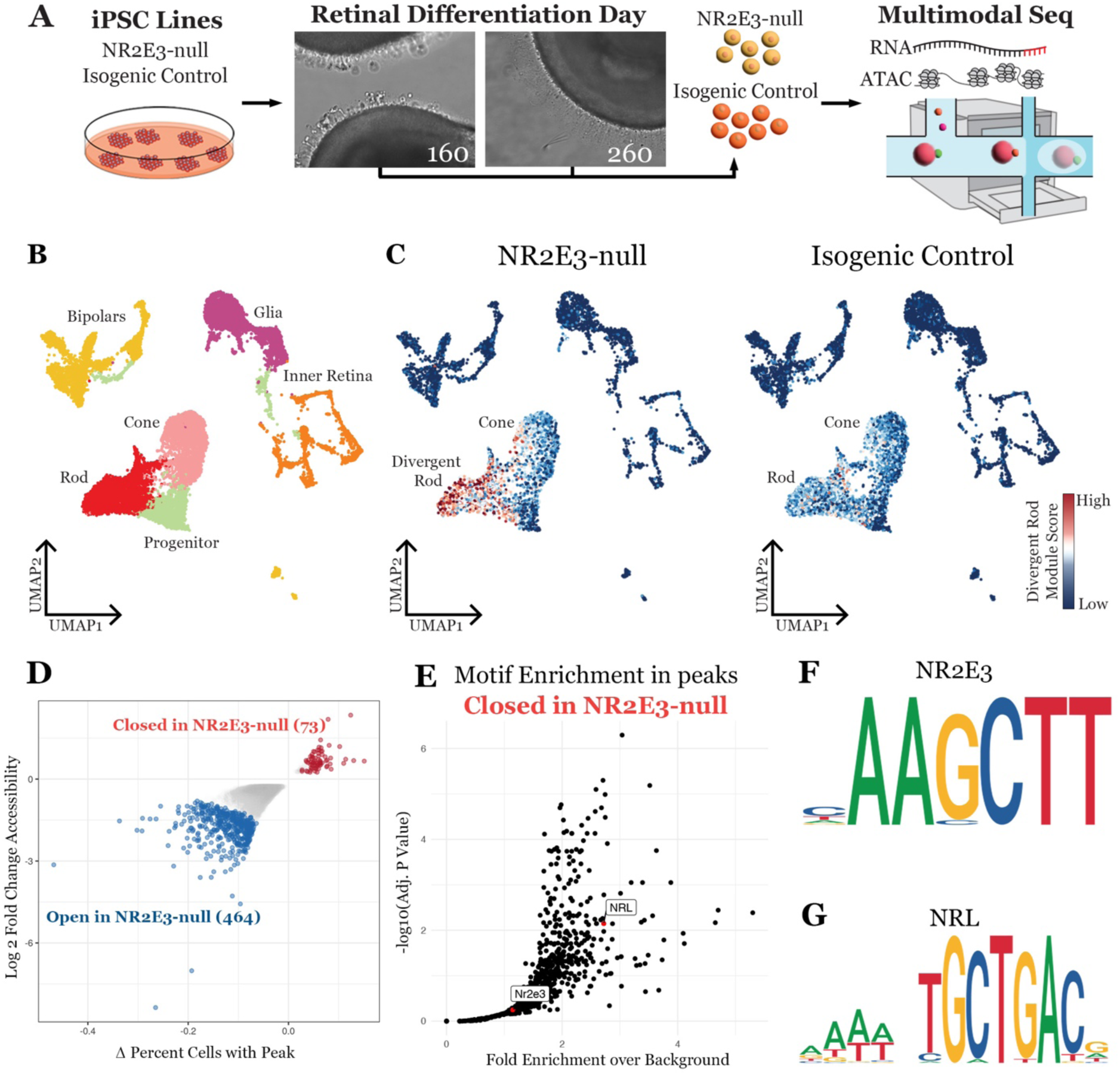
NR2E3 loss disrupts rod chromatin accessibility. **A)** Experimental schematic showing collection of nuclei from D160 and D260 retinal organoids for joint multimodal single-nucleus sequencing. **B)** Annotated WNN-UMAP projections of cells assayed by joint multimodal single-nucleus sequencing. Cells from the NR2E3-nulland isogenic control lines are shown separately. Both lines contribute to all cell type clusters. **C)** Two-dimensional projection of cells based on WNN analysis of gene expression and ATAC-seq profiles. Cells split by line (NR2E3-null and isogenic control). Cells are shaded based on Divergent Rod Gene Module score, with red indicating enrichment for Divergent Rod Module genes. **D)** Differential ATAC peak accessibility between NR2E3-null and isogenic control rods. Peaks that are more accessible in the control line (i.e. closed in the NR2E3-null rods) are shown in red while peaks that are more accessible in the NR2E3-null line are shown in blue. More peaks are accessible in NR2E3-null versus control, indicating a globally repressive role for NR2E3 in maturing rod photoreceptors. **E)** Transcription factor binding motif enrichment in peaks that are inaccessible in the NR2E3-null rods versus control. Enrichment of the NRL motif indicates reliance of NRL on NR2E3 presence for binding. **F, G)** Motif symbols for the NR2E3 and NRL-binding motifs.

### NR2E3 acts as a direct suppressor of cone-specific gene expression

Since Nr2e3 acts as a suppressor of cone photoreceptor cell fate in mouse (Cheng et al., 2006), we were interested in how loss of NR2E3 function in human iPSC-derived photoreceptors altered the chromatin accessibility around cone genes and caused their misregulation. After ATAC peaks were called, differentially accessible regions (DAR) between the NR2E3-null and isogenic control lines within the rod cluster were identified (**Figure 3D**). Notably, far more peaks were accessible in the NR2E3-null sample, indicating NR2E3’s globally repressive role (464 peaks preferentially accessible in NR2E3-null compared to control with 73 peaks preferentially inaccessible in the NR2E3-null cells). Several of these DARs were near rod- and cone-specific genes, indicating a dual role for NR2E3 in cone gene repression and rod gene activation (**Figure S4K**). We next computed enrichment of known transcription factor binding motifs within ATAC peaks that were inaccessible in NR2E3-null rod photoreceptors (**Figure 3E**). Of specific interest were the canonical NRL and NR2E3-binding motifs (**Figure 3F, G**). The NRL-binding motif (TGCTGAC) was statistically significantly enriched in the set of peaks that become inaccessible following NR2E3 loss showing that NRL binding and subsequent chromatin remodeling requires the presence of NR2E3 in at least some contexts (**Figure 3E**).

### Divergent rods misexpress cone and rod-specific phototransduction genes

We next asked how the gene expression pattern of divergent rods differed from that of normal rods and cones. A differential expression test between the most mature cluster of three lineages (rods, cones, and divergent rods, see **Figure 2B, C**) was performed in a pairwise fashion between each lineage. Remarkably, several well described functional photoreceptor genes were misexpressed in divergent rods (**Figure 4A**, highlighted in red). Cone-specific genes (e.g., *PEX5L*, *PDE6H*, *ARR3*) were upregulated in divergent rods compared to normal rods while several canonical rod genes (e.g., *NRL*, *ROM1*, *GNAT1*, *GNGT1*) were upregulated compared to normal cones. Comparison of gene expression between NR2E3-null and isogenic control cells showed that dysregulation was restricted to rod photoreceptors (**Figure S5A-N**). Normal expression patterns of rod- and cone-specific genes were confirmed using scRNAseq data from human donor retina (Voigt et al., 2021) (**Figure S5O-V**). These data indicate that these divergent rod cells retain rod identity and are not shunted to a cone fate in early differentiation; instead, they exist as mature photoreceptor cells expressing a combination of rod and cone genes.

**Figure 4.**
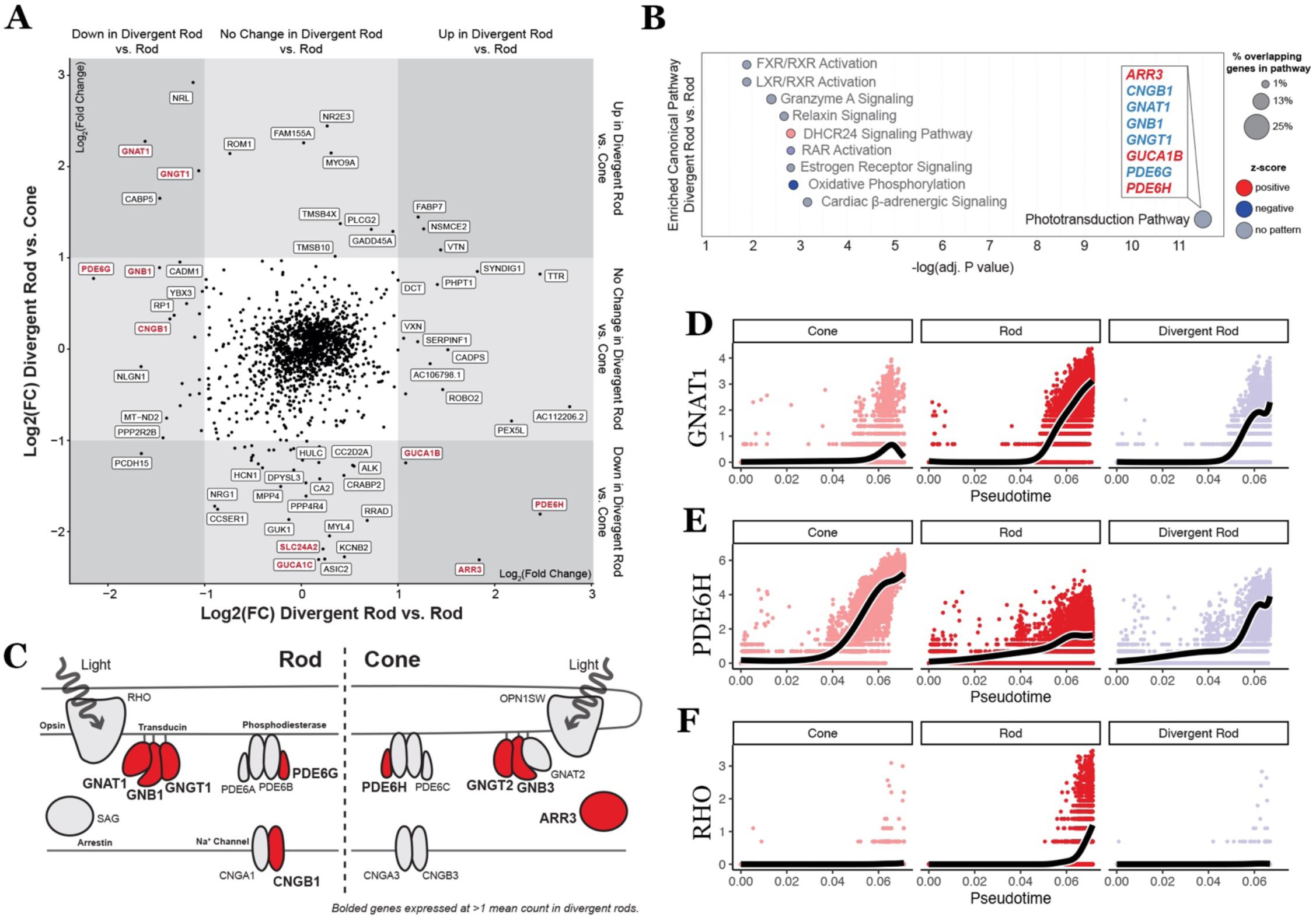
Divergent rods express a combination of rod- and cone-specific genes involved in phototransduction. **A)** Differentially expressed genes between the divergent rod and rod (x-axis) or cone (y-axis) lineages. Compared to normal rods, divergent rods upregulate several cone-specific transcripts, as well as genes involved in synaptogenesis. Compared to normal cones, divergent rods upregulate canonical rod transcripts (e.g., NRL, ROM1, GNAT1, GNGT1). Genes involved in phototransduction are highlighted in red. **B)** Pathway enrichment analysis for differentially expressed genes between divergent rod and rod clusters (the x-axis of panel **A**) shows strong dysregulation of genes involved in phototransduction. **C)** Diagram of rod-(left) and cone-(right) specific components of the phototransduction pathway. Genes expressed in divergent rods are colored and those not expressed in divergent rods are shown in gray. **D)** The rod-specific transducin component (*GNAT1*) is expressed in rod and divergent rod lineages but not in normal cones. **E)** The cone-specific phosphodiesterase *PDE6H* is expressed in the normal cone lineage and in divergent rods across the same developmental time. **F)** The rod-specific opsin *RHO* is expressed late in normal rod development but not divergent rods.

Marker genes either up- or downregulated in divergent rods versus normal cones and rods were identified (**Figure 4A**). Differentially expressed genes between divergent rods and normal rods were subjected to pathway enrichment analysis to better understand the cellular changes downstream of NR2E3 loss. The top enriched pathway in the divergent rod differentially expressed genes was “Phototransduction Pathway” (**Figure 4B**), validating the observed misexpression of genes such as *ARR3*, *PDE6H*, *GNAT1*, and *GNGT1* and *GNGT*2 in **Figure 4A**. Divergent rods express a combination of rod- and cone-specific phototransduction genes but fail to express either rod or cone opsin (**Figure 4C**). The expression of phototransduction genes along pseudotime in divergent rods compared to the normal rod and cone lineages show that misexpression follows normal timing (**Figure 4D - F**). Specifically, expression of cone genes in divergent rods temporally occurs in accordance with expression in normal cones, and expression of rod genes in divergent rods does so in accordance with normal rods. Based on these analyses, NR2E3 loss in developing photoreceptors causes misexpression of cone- and rod-specific genes involved in phototransduction, the major function of photoreceptor cells in reception and processing of visual information.

To investigate the potential functionality of divergent rods, we next examined expression of the rod-specific opsin gene, RHO. While divergent rods express *NRL* and the variant-containing *NR2E3* transcript at normal levels, no detectable *RHO* transcript was found in NR2E3-null organoids at any timepoint (**Figure 5A**). Across D160 and D260, chromatin accessibility in the *RHO* coding sequences and cis regulatory sites was greatly diminished in *NR2E3*-null rods compared to control rods (**Figure 5B**). D160 and D260 organoids were stained for RHO protein expression and a similar pattern was observed wherein no RHO protein was found in the photoreceptors of mature NR2E3-patient organoids (**Figure 5D - I**).

**Figure 5.**
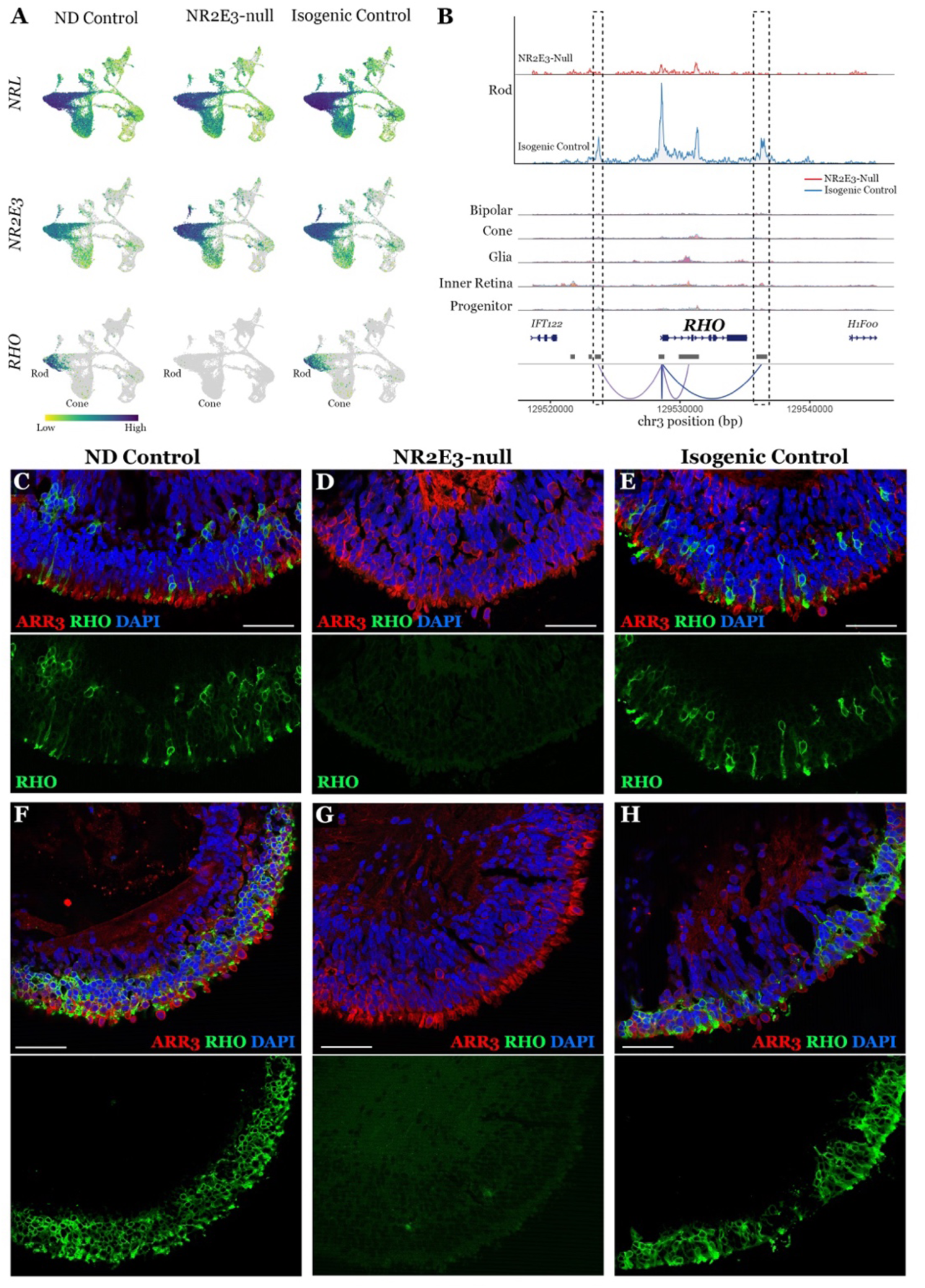
NR2E3-null rods fail to activate expression of rhodopsin. **A)** Per-cell expression levels of *NRL*, *NR2E3*, and *RHO* in ND control, NR2E3-null and isogneic control retinal organoids. All lines express *NRL* primarily in the rod cluster. All lines, including NR2E3-null, express *NR2E3* at the transcript level. The NR2E3-null line expresses no *RHO* transcript in the rod cluster or otherwise, unlike control lines. **B)** ATAC coverage tracks show chromatin accessibility at the *RHO* gene locus in control rods and NR2E3-null (i.e., divergent) rods (top) and other cell types. Linkages (bottom) indicate correlation between accessibility at a peak and transcript abundance in single cells assayed by joint multimodal sequencing. **C-E)** At D160 *RHO*-expressing photoreceptors are observed in ND control (C) and isogenic control (E) organoids but no *RHO*-expressing cells are seen in NR2E3-null organoids (D). **F-H)** By D260 *RHO* expression increases in ND control (F) and isogenic control (G) organoids but is still absent from NR2E3-null organoids (H). Scalebar = 50um.

### NR2E3 suppresses cone-specific gene expression in rods

To confirm that divergent rods genuinely co-express both rod and cone genes, we subsetted the rod, cone, and divergent rod populations and plotted cells along two axes for the canonical rod transducin component G protein subunit alpha transducin 1 (*GNAT1*) and the canonical cone phosphodiesterase 6H (*PDE6H*) (**Figure 6A-C**). In both non-diseased (**Figure 6A**) and isogenic (**Figure 6C**) control samples, few cells co-expressed rod- and cone-specific genes, with most photoreceptors exclusively expressing either *PDE6H* or *GNAT1*. However, divergent rods found in the NR2E3-null organoids largely co-expressed both genes (**Figure 6B**). Such co-expression of *GNAT1* and *PDE6H* was further visualized at the protein level in mature organoids (**Figure 6D-F**). Colocalization of GNAT1 and PDE6H protein in photoreceptor outer segments was never observed in the control lines (**Figure 6D, F**) but was commonly observed in the NR2E3-null organoids (**Figure 6E**). These findings were confirmed by scRNAseq of late stage (D260) organoids (**Figure S6**).

**Figure 6.**
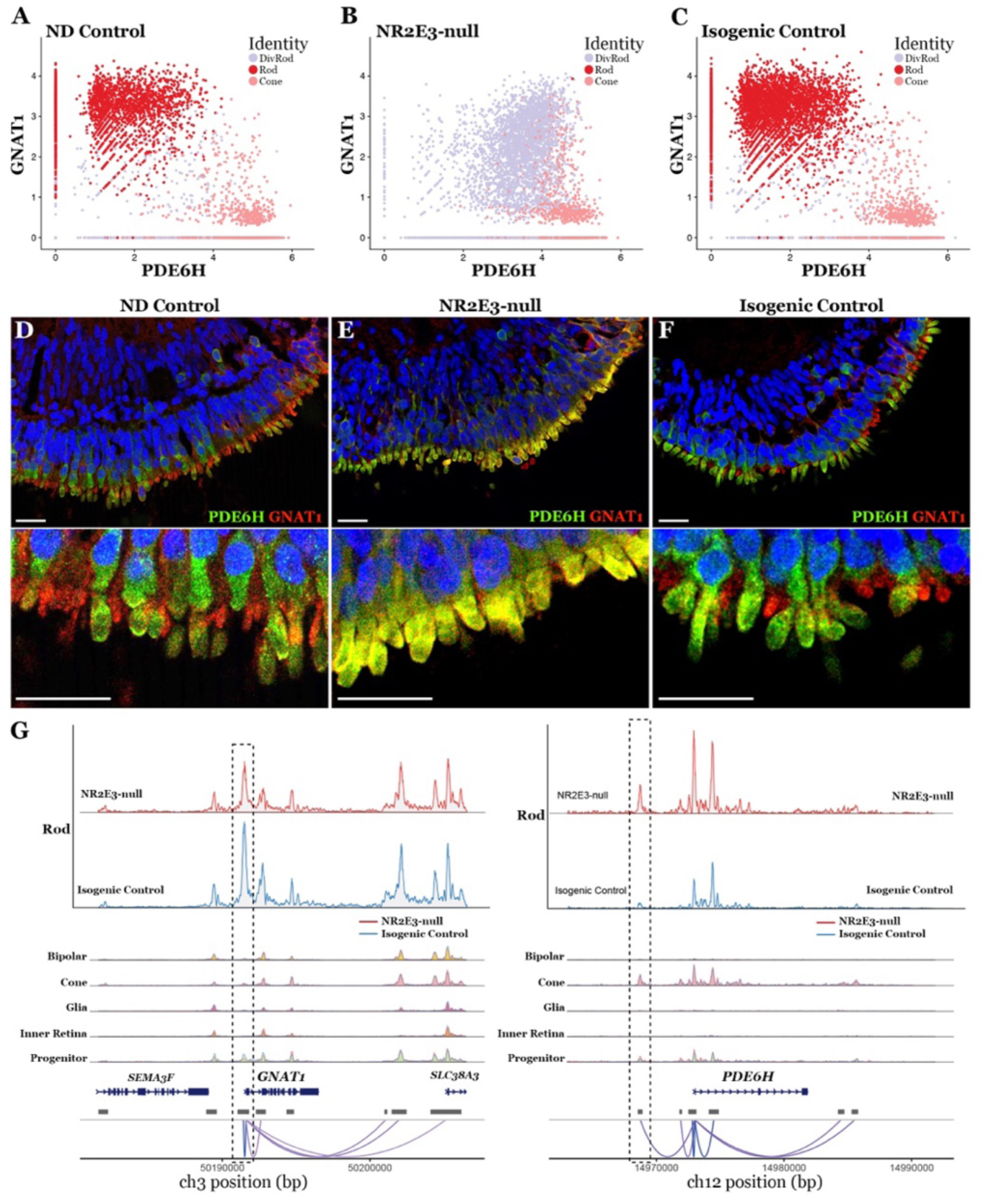
NR2E3 is required for repression of cone-specific phototransduction genes. **A)** Cells of the photoreceptor lineage from the ND control line are plotted based on expression level of PDE6H (x-axis) and GNAT1 (y-axis). Photoreceptor cells segregate by class with rods expressing GNAT1 and cones expressing PDE6H. **B)** Divergent rods co-express PDE6H and GNAT1 at high levels. No cells are observed to express only GNAT1, indicating lack of a normal rod population. PDE6H-expressing cone population appears normal. **C)** The isogenic control line restores normal segregation of photoreceptor classes. Clusters resemble those of the ND control in A. **D)** Segregation of expression of GNAT1 (red) and PDE6H (green) into rods and cones is observed in D260 ND control retinal organoids. **E)** Photoreceptors from the NR2E3-null organoids exhibit colocalization of GNAT1 and PDE6H protein. **F)** Segregation of expression is restored in isogenic control organoids. **G)** ATAC coverage tracks for the rod cluster of organoids (D160 and D260 combined) are shown at the top for NR2E3-null and isogenic control samples. Other cell type tracks show all samples combined. The *GNAT1* locus is shown on the left. A peak linked to expression of *GNAT1* is shown boxed. This peak is accessible in both NR2E3-null and isogenic control rods. On the right, the *PDE6H* locus is shown. NR2E3-null rods show accessibility at a peak normally accessibly only in cones. This peak is linked to expression of *PDE6H*, indicating a de-repression caused by NR2E3 loss leads to inappropriate *PDE6H* expression. Scalebar = 50μm.

GNAT1 is a canonically rod-specific component of the transducin complex. Accessibility at three nearby ATAC peaks are found to be significantly correlated with expression of the GNAT1 gene (**Figure 6G**, peak to gene links indicated at bottom). Of these, one peak (left box, **Figure 6G**) is accessible only in rods and photoreceptor progenitors. This rod-specific peak is observed in both NR2E3-null and isogenic control lines, indicating that loss of NR2E3 activity has no deleterious effect on rod-specific chromatin remodeling or transcription of GNAT1 (as previously observed in **Figure 6B & E**).

Loss of transcriptional repression of a cone-specific gene at the chromatin level following loss of NR2E3 is observed in the regulatory region surrounding *PDE6H* (**Figure 6G**). A cone-specific peak linked to expression of PDE6H is observed in NR2E3-null rods (**Figure 6G**, right box), indicating a failure to repress expression downstream of NR2E3 loss of this cone-specific member of the phosphodiesterase complex. The same peak in this presumptive enhancer of expression is observed in NR2E3-null rods and all cones (**Figure 6G**, right box). Together, these results show how NR2E3-null rods fail to suppress cone genes involved in phototransduction.

### NR2E3-null divergent rods are transcriptionally distinct from NRL-null cods

Enhanced S-cone syndrome is most often caused by mutation in *NR2E3* (Yzer, 2013). However, rare cases of ESCS are known to be caused by mutations in the genetically upstream rod photoreceptor-specific transcription factor gene *NRL* (Kallman et al., 2020). A previous study of human retinal organoids lacking NRL activity described the presence of hybrid cone/rod photoreceptor cells termed “cods”, described earlier in the Nrl^-/-^ mouse (Kallman *et al*., 2020; Mears *et al*., 2001). As NRL is the inducer of NR2E3 expression in normal rod development (Cheng et al., 2006), cods lack NR2E3 expression (Kallman et al., 2020). We therefore asked how divergent rods differed at the transcriptome-level from cods to describe the specific contribution of NR2E3 to rod development genetically and temporally downstream of NRL. We generated scRNAseq profiles of D260 organoids (**Figure S6**), which showed persistence of divergent rods at a comparable timepoint to that studied by (Kallman *et al*., 2020).Our data were integrated with that of Kallman and the divergent rod and cod populations were annotated (see methods) and highlighted (**Figure 7A**). Differential expression analysis was performed between divergent rods and normal rods from the current study and between cods and normal rods from the Kallman study to remove confounding effects introduced by differences in the differentiation protocol used by each study. The fold change of each analysis for each gene is plotted (**Figure 7B**). Remarkably, several genes are dysregulated only in cods or divergent rods respectively. That is, cods cluster more closely to normal cones, while divergent rods cluster more closely to normal rods, **Figure 7A-B**. This indicates that NR2E3 and NRL regulate partially exclusive subsets of genes essential to rod development and function.

**Figure 7.**
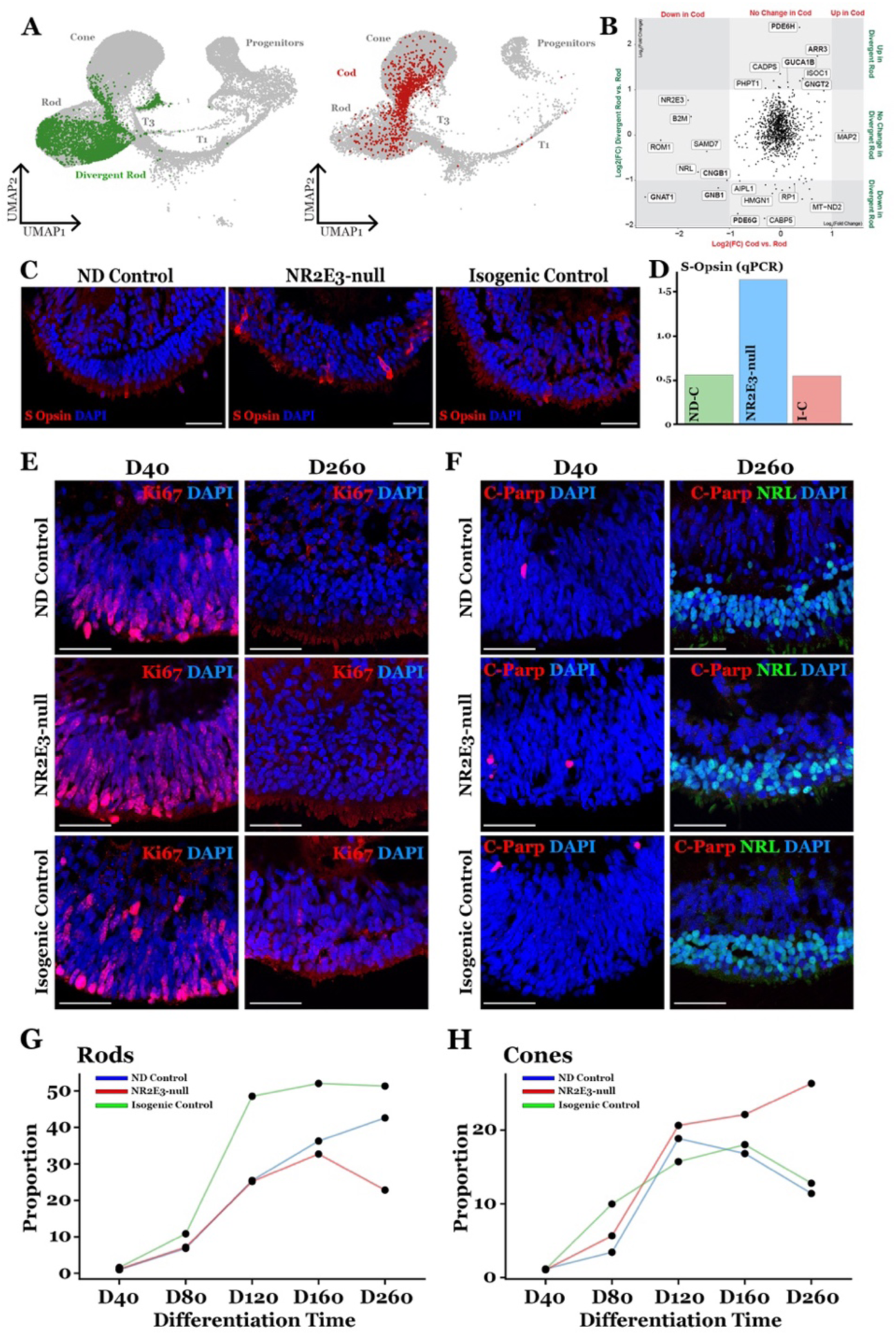
Divergent rod fate in the context of Enhanced S-Cone Syndrome. **A)** D40 – D260 data from the photoreceptor lineage of the current study integrated with the same cell types from Kallman et al. Cells are shown split by study and projected in 2D space using UMAP embeddings. Divergent rods are colored green and NRL-null cods are colored red. **B)** Differential expression analysis between pathologic and normal rods from each study. Highlighted genes are significantly dysregulated in NRL- and/or NR2E3-null cells compared to control rods. **C)** D260 retinal organoids from the current study stained for S-opsin (OPN1SW). NR2E3-null organoids display a modest increase in the proportion of S-opsin-expressing cells. **D)** RT-qPCR shows an increase in OPN1SW transcript in NR2E3-null organoids compared to controls. **E)** The marker of proliferation Ki67 is expressed in D40 organoids and lost at D260 in all three lines, indicate no prolongation of proliferation in NR2E3-null organoids. **F)** No increase in cleaved PARP is observed in NR2E3-null organoids, indicating no increased apoptosis. **G-H)** Between D160 and D260 the rod proportion of NR2E3-null organoids decreases while the cone proportion increases. The opposite trend is observed in controls. Scalebar = 50μm.

In addition to a lack of normal rod photoreceptor cell function, ESCS is characterized by an exaggerated retinal response to short wavelength light. Previous studies in NRL-null organoids (Kallman et al., 2020) and post-mortem examinations of NR2E3-associated ESCS eyes (Milam et al., 2002) have shown an increased number of S-opsin expressing cells in the ESCS photoreceptor cell layer. However, there are drastic differences in the observed magnitude of fate conversion of rods to S-cones in NRL-null animal models versus NR2E3-null models and patient observations. NRL-null organoids exhibited a complete conversion, with seemingly all rods becoming OPN1SW-expressing cods early in development. Milam, et al., showed that in NR2E3 patient retinas, the number of cone photoreceptors was only increased two-fold, with the vast majority of those expressing OPN1SW. These findings leave open the possibility of divergent rods persisting in the adult NR2E3 patient retina. In fact, we observed only a modest increase in S-opsin expressing cells and *OPN1SW* transcript in NR2E3-null organoids (**Figure 7C, D**) and no change in the amount of ML-opsin expressing cells (**Figure S7G - I**). We integrated our scRNAseq data from D40 to D260 of differentiation to understand the late fate of divergent rods. By D260, no early or intermediate progenitors existed (**Figure S7**), and the proportion of other major cell types was equivalent between lines **(Figure S7**). Examining the rod cluster, we observed a decrease in the proportion of rods between D160 and D260 only in the NR2E3-null line (**Figure 7G**). We observed the opposite trend in the cone cluster (**Figure 7H)**, where the proportion of cones in the NR2E3-nulls organoids increased between D160 and D260, decreasing in the control lines. This is not the result of proliferation of progenitor cells or death of a certain population of cell because there is no difference in ki67 (proliferation marker) or cleaved PARP (apoptosis marker) at D40 or D260 (**Figure 7E, F**). These data suggest that a subset of divergent rods may give rise to late-born cone photoreceptors in NR2E3-null retinal organoids.

## DISCUSSION

Much of the cellular biology underlying human retinal disease must be inferred from clinical imaging studies, animal models, or post-mortem case studies. This is primarily due to the relative inaccessibility of retina in living patients for research studies compared with easily biopsied tissues such as blood. As such, the precise downstream effects of loss of developmental genes in the human retina is largely unknown. In this study, we used patient-derived iPSC-based modeling to capture developing human retinal cells with a clinically relevant mutation. We show that sampling developing organoids at multiple timepoints using scRNAseq allows precise identification of the immediate consequences of NR2E3 loss following rod photoreceptor commitment. We demonstrate that NR2E3 loss in human retinal cells specifically causes misregulation of several rod- and cone-specific phototransduction genes, including loss of *RHO* expression. Using joint multimodal single-cell RNA and ATAC sequencine, we identify the putative cis-regulatory sites of the misexpressed phototransduction genes and show how loss of NR2E3 alters chromatin accessibility in both activating and repressive roles. Interestingly, loss of Nr2e3 in mouse does not have the same effect. Specifically, deletion of Nr2e3 expression in mouse does not appear to alter rhodopsin expression. Together, we show how an *in vitro* model of human retinal development can be used to parse the species-specific roles of a core developmental transcription factor with implications for inherited retinal disease.

### Characterization of divergent rods

In this study, we identified a novel population of rod photoreceptor cells unique to the NR2E3-null disease state. Importantly, NR2E3-null retinal progenitors differentiate normally until a late timepoint, after the induction of NRL specifies rod photoreceptors. Because of this, prior to D120, divergent rod photoreceptor cells are indistinguishable from normal rods and earlier progenitors. These data indicate that NR2E3 loss in human cells does not delay progenitor commitment nor shunt presumptive rods into a cone fate during early development, counter to hypotheses based on clinical imaging and mouse models. Instead, the primary defect of NR2E3-null divergent rods seems to involve late genes involved in maturation and rod function, rather than commitment. Of note, this is dissimilar to the defect observed by Kallman and colleagues in the NRL-null human retinal organoids (Kallman et al., 2020), wherein cods appeared to be in a state of arrested commitment, expressing markers of T3 photoreceptor progenitors such as *FABP7* and *ISOC1* (described in human fetal retina and retinal organoids (Sridhar et al., 2020)).

### NR2E3/NRL cooperation in rod development

Rhodopsin (*RHO*) is the light-sensitive opsin protein that allows rod photoreceptors to detect and trigger the signals required to enable vision in dim light. Rhodopsin is expressed exclusively in rods following early differentiation from uncommitted photoreceptor progenitors. Understanding how and when upstream factors control *RHO* expression is essential to addressing photoreceptor dysfunction in the disease state. Expression of *RHO* is thought to be under the control of the rod-specific transcription factors NRL and NR2E3; however, several lines of evidence imply major regulatory differences between human and murine *RHO* regulation. As indicated above, NR2E3-deficient mice express RHO in rod photoreceptors, as do NRL-deficient animals expressing NR2E3 under other rod-specific promoters (Cheng et al., 2006), which suggests a compensatory relationship between NR2E3 and NRL wherein either is sufficient to drive RHO expression in mouse. In human retinal organoids lacking NRL, neither NRL or NR2E3 are functional, and RHO is not expressed (Kallman et al., 2020). Here, we show that even in rod photoreceptors with normal levels of functional NRL, loss of NR2E3 is sufficient to prevent *RHO* expression.

### Supernumerary S-cones in NRL-null retina

ESCS is characterized by an increased retinal sensitivity to short wavelength light, mediated by S-cones, and a loss of sensitivity to dim light, mediated by rod photoreceptors. Due to the known role of NR2E3 in rod photoreceptor fate specification, supernumerary S-cone formation in lieu of rod formation has been proposed as the explanation for cone hypersensitivity. Post-mortem study of ESCS patient retinas revealed a slight increase in the proportion of cones, approximately two-fold greater than normal, with most of these expressing S-opsin (Milam et al., 2002). This is similar to what we observed in the current study (**Figure 7**) but is contrary to what is observed following loss of *NRL* expression (Kallman, 2020). Interestingly, the increase in S-cone number in our study was not readily apparent until 260 days of differentiation, which is quite late in development. With an increased proportion of blue cones, we observed a concomitant decrease in the proportion of divergent rods. Since there was no increase in the rate of apoptosis to suggest divergent rod cell death, we hypothesized that under prolonged cell culture divergent rods may transdifferentiate into S-cones. This could occur by either direct conversion or reversion to an earlier developmental state followed by progression down a blue cone developmental pathway. To evaluate this hypothesis, further experimentation will be required to determine if a subpopulation of *NRL*-expressing divergent rods are able to silence rod gene expression and revert to a bona fide blue cone cell state.

### Conclusion

In summary, in this study we demonstrate the power of a combined patient derived induced pluripotent stem cell, CRISPR-based genome editing, and single-cell sequencing strategy to elucidate the role of the transcription factor NR2E3 in human retinal development and disease. Remarkably, we demonstrate that loss of the transcription factor, which is essential for rod photoreceptor cell fate commitment, has a very different outcome in human than it does in rodent. These differences are critical for understanding how the human retina develops.

### Limitations of study

Modeling retinal development with patient-derived stem cells and retinal organoids offers a unique and clinically relevant view of the molecular changes downstream of transcription factor loss. However, the organoid model is subject to several limitations. While retinal organoids produce the neuroectoderm-derived cell types of the retina, other potentially relevant cells are not present. These include immune cells such as microglia that may play a role in pruning divergent rod cells in vivo. That said, NR2E3-associated ESCS is often stationary, for instance the patient from which the iPSC line in this study was generated has a remarkably stable visual acuity and field. This suggests that divergent rods may persist late into adulthood. While we attempted to infer functionality of divergent rods based on expression of key phototransduction genes at the protein and transcript level, further work will be needed to determine what, if any, electrical response to light is present in such cells. Finally, while we observed an increase in the proportion of S-cone concomitant with a decrease in divergent rod proportion between D160 and D260 with no obvious changes to proliferation or cell death, further work will be required to lineage trace developing divergent rods and determine their fate in developing organoids.

## METHODS

### Patient-derived iPSC generation and validation

This study was approved by the Institutional Review Board of the University of Iowa (project approval #200202022) and adhered to the tenets set forth in the Declaration of Helsinki. Patient iPSCs were generated from an individual with no disease and an individual with molecularly confirmed enhanced S cone syndrome (ESCS) as described previously (Bohrer *et al*., 2019; Wiley et al., 2016). The disease-causing mutation in NR2E3, c.119-2A>C, was corrected via CRISPR-mediated homology dependent repair in patient-derived iPSCs as described previously (Bohrer *et al*., 2019). Following generation and clonal expansion, iPSCs were karyotyped in metaphase by the Shivanand R. Patil Cytogenetics and Molecular Laboratory at the University of Iowa using Leica Microsystems metaphase scanning platform and CytoVision version 7.7 software as previously demonstrated (Bohrer *et al*., 2019). Briefly, cells were grown *in vitro* and arrested at metaphase with colcemid. Chromosomes were stained by the G-banding method, counted, and structurally evaluated for the presence or absence of detectable rearrangements. At least 20 cells were analyzed for each iPSC line.

### Retinal organoid differentiation

Retinal differentiation was performed as described previously with minor modifications (Bohrer *et al*., 2019). Briefly, iPSCs were cultured on laminin 521 coated plates in E8 medium. Embryoid bodies (EBs) were lifted with ReLeSR (STEMCELL Technologies, Cambridge, MA) and transitioned from E8 to neural induction medium (NIM-DMEM/F12 (1:1), 1% N2 supplement, 1% non-essential amino acids, 1% Glutamax (Thermo Fisher Scientific), 2 μg/mL heparin (Sigma-Aldrich, St. Louis MO) and Primocin (InvivoGen, San Diego, CA)) over a four-day period. On day 6, NIM was supplemented with 1.5 nM BMP4 (R&D Systems, Minneapolis, MN). On day 7, EBs were adhered to Matrigel coated plates (Corning). BMP4 was gradually transitioned out of the NIM over seven days. On day 16, the media was changed to retinal differentiation medium (RDM - DMEM/F12 (3:1), 2% B27 supplement (Thermo Fisher Scientific), 1% non-essential amino acids, 1% Glutamax and 0.2% Primocin). On day 25-30 the entire EB outgrowth was mechanically lifted using a cell scraper and transferred to ultra-low attachment flasks in 3D-RDM (RDM plus 10% fetal bovine serum (FBS)); Thermo Fisher Scientific), 100 µM taurine (Sigma-Aldrich), 1:1000 chemically defined lipid concentrate (Thermo Fisher Scientific), and 1 µM all-trans retinoic acid (until day 100; Sigma-Aldrich). The cells were fed three times per week with 3D-RDM until harvest.

### Immunocytochemistry

Organoids were fixed with 4% paraformaldehyde for 30-60 minutes at room temperature and equilibrated to 15% sucrose in PBS, followed by 30% sucrose. Organoids were cryopreserved in 50:50 solution of 30% sucrose/PBS: tissue freezing medium (Electron Microscopy Sciences, Hatfield, PA) and cryosectioned (15 μm). Sections were blocked with 5% normal donkey serum, 3% bovine serum albumin (BSA), and 0.1% triton-x and stained overnight with the primary antibodies listed in **Table S1**. The following secondary antibodies (Thermo Fisher Scientific) were incubated for 1 hour: donkey anti-goat 488 (Cat# R37118), donkey anti-mouse 488 (Cat# A21202), donkey anti-sheep 647 (Cat#A21448) and donkey anti-rabbit 647 (Cat# A31573). Cell nuclei were counterstained using DAPI (Thermo Fisher Scientific; Cat# 62248). Images were acquired using a Leica TCS SPE upright confocal microscope system (Leica Microsystems, Wetzlar, Germany)

### RT-qPCR

RNA was isolated from snap-frozen retinal organoids using the NucleoSpin RNA kit (Macherey-Nagel) following manufacturer’s instructions. cDNA was synthesized using the High Capacity cDNA Reverse Transcription Kit (Applied Biosystems) with 120ng RNA as input. Probe-based qPCR was carried out for the S-opsin transcript (*OPN1SW*) using predesigned or custom assays (Integrated DNA Technologies) and normalized to expression of *HPRT*. Reactions were set up using TaqMan Advanced Fast Master Mix (Invitrogen) and run on the QuantStudio3 instrument (Invitrogen). Based on input of RNA to cDNA synthesis, the equivalent of 1ng of cDNA was used per qPCR reaction.

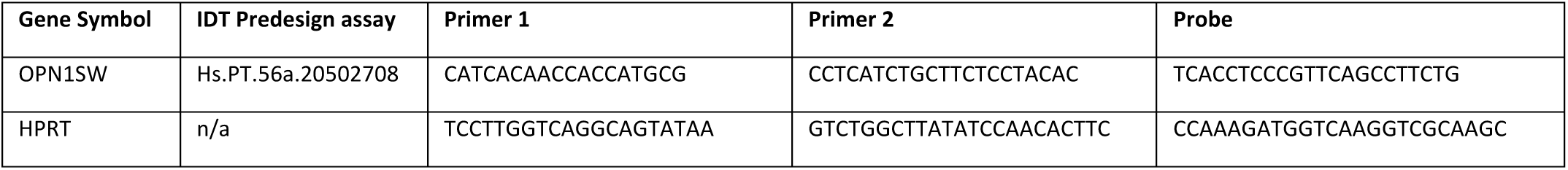

### Organoid dissociation for single-cell RNAseq

Samples were collected at differentiation day 0, 40, 80, 120, 160, and 260 and processed for single-cell gene expression profiling. Day 0 (undifferentiated iPSC) samples were lifted with TrypLE (Gibco) and washed once with dPBS. For Day 40-260 samples, approximately ten organoids displaying morphology of successful retinal differentiation were selected for each of the three lines. Organoids were allowed to settle by gravity in a 1.5mL tube and culture medium was removed. Organoids were dissociated in a solution of 20U/mL papain (Worthington) and 120U/mL DNase (Worthington) in Earle’s balanced salt solution (EBSS) (Worthington). Samples were incubated in 300µL of above papain solution at 37°C with continuous shaking (500rpm) until all organoids were completely dissociated (approximately 1 hour). Samples were triturated with a pipette every 15 minutes. Following dissociation, cells were pelleted at 500 x g for 5 minutes and resuspended in 8µg/mL recombinant albumin (New England Biolabs) in dPBS-/-. Cells were passed through a 70µm filter to encapsulation with the Chromium Controller instrument (10X Genomics). Approximately 8,000 cells were targeted for encapsulation per sample.

### Nucleus isolation for single-nucleus multimodal sequencing

Nuclei were isolated from D160 and D260 organoids cultured in an independent batch from those used in the scRNAseq study above. Nuclei were isolated following a protocol based on 10X Genomics’ demonstrated protocol CG000366. Briefly, approximately ten organoids were selected as described above. Medium was aspirated and 500µL of chilled 0.1X Lysis Buffer (Tris-HCl, NaCl, MgCl_2_, Tween-20, NP40, Digitonin, BSA, DTT, RNase Inhibitor) was added to each sample. Samples were homogenized with 15 strokes of a sterile pestle in a 1.5mL tube on ice. Samples were then incubated on ice for 5 minutes. Following incubation, samples were mixed with a P1000 pipette 10 times on ice and incubated for 8.5 minutes on ice. 500µL of Wash Buffer (Tris-HCl, NaCl, MgCl_2_, Tween-20, BSA, DTT, RNase inhibitor) was added and lysed cells were mixed 5 times with pipetting. Cells were pelleted at 500 x g for 5 minutes in a refrigerated (4°C) centrifuge and resuspended in 500µL chilled wash buffer. This process was repeated for three total washes. Nuclei were resuspended in Diluted Nuclei Buffer (Nuclei Buffer (10X Genomics), DTT, RNase inhibitor) and passed through a 70µm strainer. Nuclei were counted on a hemocytometer with DAPI to visualize intact nuclei. Approximately 9,000 nuclei were targeted for encapsulation per sample using the Chromium X instrument (10X Genomics).

### Single-cell gene expression library preparation and sequencing

Single cells were partitioned and barcoded with the Chromium Controller instrument (10X Genomics) and Single Cell 3’ Reagent (v3.1 chemistry) kit (10X Genomics) according to the manufacturer’s specifications with no modification (Rev C). Final libraries were quantified using the Qubit dsDNA HS Assay Kit (Life Technologies) and diluted to 3ng/µL in buffer EB (Qiagen). Library quality was confirmed using the Bioanalyzer High Sensitivity DNA Assay (Agilent) prior to sequencing.

### Single-nucleus multimodal library preparation and sequencing

Nuclei were processed following the 10X Genomics Chromium Next GEM Single Cell Multiome ATAC + Gene Expression (Rev. E) without modification. Final libraries were quantified using the Qubit dsDNA HS Assay Kit and diluted to 3ng/µL. Library quality was checked using the Bioanalyzer or Tapestation (Agilent).

### Single-cell gene expression data integration and processing

scRNA libraries were pooled and sequenced using the NovaSeq 6000 instrument (Illumina) generating 100-bp paired end reads. Sequencing was performed by the Genomics Division of the Iowa Institute of Human Genetics. FASTQ files were generated from base calls with the bcl2fastq software (Illumina), and reads were mapped to the pre-built GRCh38 reference (refdata-gex-GRCh38-2020-A) with Cell Ranger v7.0.0 (10X Genomics) using the ‘count’ function with the following parameters: --expect-cells=8000 -- localcores=56. Only cells passing the default Cell Ranger call were analyzed further. Differentiation day 40-160 samples were integrated using canonical correlation analysis (CCA) in Seurat v4.0.3 (40). Only cells with between 1,000 and 7,000 unique genes (features) were included in the analysis. Only cells with < 10% of reads mapping to mtDNA-encoded genes and < 20% of reading mapping to ribosomal genes were included. Counts data were normalized using the NormalizeData function (Seurat) with the following parameters: normalization.method = “LogNormalize”, scale.factor = 10000. 2000 variable features were identified with the FindVariableFeatures function using the vst selection method. Integration anchors were identified using the FindIntegrationAnchors function using 25 dimensions. An assay “Integrated” was generated for the 2000 variable features using the IntegrateData function using 25 dimensions. The integrated data were then used in principal component analysis (PCA).

### Single-nucleus multimodal data integration and processing

Single-nucleus multimodal libraries were sequenced using the NovaSeq 6000 instrument (Illumina). Sequencing was performed by the Genomics Division of the Iowa Institute of Human Genetics. FASTQ files were generated from base calls with the bcl2fastq software (Illumina). Reads were mapped to the pre-built GRCh38 reference (GRCh38-2020-A-2.0.0, 10X Genomics) using Cell Ranger ARC (v.2.0.0, 10X Genomics) with default parameters. Resulting cell-by-peak and cell-by-gene matrices (ATAC and Gene Expression assays respectively) from the four samples were integrated separately using latent semantic indexing (ATAC) and canonical correlation analysis (gene expression).

### Dimensionality reduction with UMAP and cell type annotation

30 principal components were identified out of the D40-D160 integrated dataset described above using the RunPCA function (Seurat) (Stuart et al., 2019). Uniform manifold approximation and projection (UMAP) was performed using the RunUMAP function using 25 principal components. Cells were annotated with the FindTransferAnchors and TransferData functions using 30 principal components and data from (Sridhar et al., 2020) as a reference. The photoreceptor cluster was manually refined to Rod and Cone photoreceptor cell classes based on expression of several canonical marker genes (e.g., *ARR3*, *GNGT1*, *RCVRN*, etc.).

### Dimensionality reduction of multimodal data with WNN-UMAP and cell type annotation

Only cells passing both gene expression and ATAC assay quality control metrics were used in downstream analysis. A weighted nearest neighbor (WNN) graph was constructed based on a weighted combination of either sequencing modality (GEX and ATAC). From the GEX data, dimensions 1:30 of the PCA reduction from CCA were used. From the ATAC data, dimensions 2:50 of the LSI were was used. UMAP was performed on the resulting WNN graph. Clusters were identified based on the weighted shared nearest neighbor graph using the SLM algorithm with a resolution of 0.5 using the FindClusters function in Seurat. Cluster identity (i.e., cell type) was assigned based on expression (from the RNA assay) of cell type-specific marker genes.

### Dimensionality reduction and cell type annotation with PHATE

Using the annotations described above from the differentiation day 40-160 dataset, cells within the photoreceptor developmental lineage were identified (i.e., “Progenitors”, “T1”, “T3”, “Rod”, and “Cone”). SCTransform (Hafemeister and Satija, 2019) was performed on this subset to scale and normalize the counts. PHATE (Moon et al., 2019) was run on the SCT data using the following parameters: knn=6, decay=50, t=100. Next, clustering was performed on the PHATE-derived embedding. Neighbors were identified using the FindNeighbors function (Seurat) using dimensions 1 and 2 of the PHATE reduction. Clusters were identified using the FindClusters function (Seurat) with the following parameters: resolution=0.5, algorithm=3. The resulting clusters were combined and annotated based on timepoint and cell type annotation from PCA-based clustering in the previous section.

### Trajectory construction with Slingshot

Slingshot (Street et al., 2018) was used to infer trajectories through the PHATE reduction of the photoreceptor lineage and assign pseudotime values to cells based on principal curves through identified trajectories. The slingshot function was run taking PHATE embeddings and PHATE-derived cluster labels as input”. The starting (Progenitor) and end (Cone, Rod, Divergent Rod) clusters were given, and the following parameters were used: extend = “n”, stretch = 0.1, thresh = 0.1, approx_points = 150. Three trajectories were identified. Curves were drawn based on these trajectories from which pseudotime values were derived and given to each cell.

### Differential gene expression analysis

The FindMarkers function of Seurat was used to identify differentially expressed genes between divergent rods and rods and between divergent rods and cones. Only features (genes) with a mean count of at least one across all cells in the photoreceptor lineage (Progenitors, T1, T3, Rod, Cone) were used. Genes with a Log2(Fold Change) of at least 1 and an adjusted P value below 0.05 were considered significantly differentially expressed. These genes were used to identify enriched pathway using the Ingenuity Pathway Analysis application.

### Transcription factor binding motif enrichment analysis

ATAC peaks to be used in motif enrichment analysis were first called using MACS2 (Feng et al., 2012; Zhang et al., 2008) with the CallPeaks function in Signac. Cells were grouped by cell type as described above. To identify transcription factor binding motifs enriched in differentially accessible regions in rods and divergent rods, the FindMarkers function was used with the following parameters: subset.ident = “Rod”, only.pos = FALSE, test.use = “LR”, latent.vars = “nCount_macs2_peaks”. Differentially accessible regions were filtered for a p < 0.05. Enriched motifs were identified in differentially accessible regions using the FindMotifs function, with a set of 50000 GC-matched peaks accessible in the same cell type used as background. Enrichment over background and significance of motifs were plotted, highlighting the motifs of interest (i.e. NRL and NR2E3).

### Peak-to-gene linkage analysis

Peak-to-gene linkages were identified using the strategy described in SHARE-seq (Ma et al., 2020) and implemented with Seurat/Signac (Hao et al., 2022). Links were computed between genes known to be involved in phototransduction (i.e., GRK, GRK7, RCVRN, RHO, SAG, GNAT1, GNAT2, GNB1, GNGT1, RGS9, PDE6A, PDE6G, PDE6B, PDE6H, GUCY2F, GUCY2D, GUCA1A, GUCA1B, GUCA1C, SLC24A1, CALML6, CALML5, CALM1, CALM2, CALM3, CALML3, CALML4, CNGB1, CNGA1) and proximal ATAC peaks as identified by MACS2 (above). Links were calculated using the LinkPeaks function (Signac) with the following custom parameter: distance = 10e+05. Links were visualized using the CoveragePlot function (Signac).

### Integration and comparison with previously published scRNAseq data

Single cell sequencing data from NRL-null and control retinal organoids were accessed from GSE143669 (Kallman *et al*., 2020). Data from the current study (D40-D260) and data from Kallman et al. were integrated using CCA. Counts were normalized using LogNormalize in the NormalizeData function in Seurat and 2,000 variable features were identified using the vst selection method. Integration anchors were identified using the first 25 principal components. UMAP was run using the first 25 principal dimensions of the integrated object. Cell types were annotated using the FindTransferAnchors function using the D40-D260 dataset from the current study as the reference.

For this analysis, divergent rods were defined using the following criteria:

1. from the current study
2. from D160 or D260 timepoint
3. predicted annotation as rod from the above label transfer
4. from the NR2E3-null line.

Cods (i.e., NRL-null hybrid photoreceptors) were defined using the following criteria:

1. from the Kallman et al. study
2. rod gene module score < 2
3. cone gene module score < 2
4. OPN1SW counts > 0.5
5. from the NRL-null line (NRL_L75P)
6. predicted annotation as either rod, cone, T1, T3, or PRC/Photoreceptor from above label transfer.

Differentially expressed genes between cods and rods from the Kallman dataset and between divergent rods and rods from the current dataset were identified using the FindMarkers function in Seurat with min.pct = 0 and logfc.threshold = 0. To generate the plot in **Figure 7B**, the gene list was filtered to only genes expressed in at least 10% of rods in both datasets.

## ACKNOWLEDGEMENTS

This work was supported by the National Institutes of Health (T32GM008629, T32GM139776, F30EY034009, F30EY031923, R01EY033308, and R01EY033331). The authors would like to acknowledge the extraordinary generosity of the patients whose donation of material made this study possible. We would also like to thank the members of the Institute for Vision Research for their helpful conversation and advice. Data presented herein were obtained at the Genomics Division of the Iowa Institute of Human Genetics which is supported, in part, by the University of Iowa Carver College of Medicine. We would like to acknowledge specifically the advice and support of M. Boes, G. Hauser, K. Knudtson, A. Sheehan, and E. Snir. Colors used in plots were based on www.colorbrewer2.org by Cynthia A. Brewer, Department of Geography, Pennsylvania State University.

## AUTHOR CONTRIBUTIONS

Conceptualization, N.K.M., L.R.B., and B.A.T.; Methodology, N.K.M., L.R.B., and A.P.V., Formal Analysis, N.K.M., L.R.B., and A.P.V.; Investigation, N.K.M., L.R.B., L.P.L., and A.W., Resources, E.M.S., and B.A.T.; Writing – Original Draft, N.K.M. and L.R.B.; Writing – Review & Editing, A.P.V., L.P.L., A.W., R.F.M., E.M.S., and B.A.T.; Funding Acquisition, N.K.M., A.P.V., R.F.M., E.M.S., and B.A.T.; Supervision, R.F.M. and B.A.T.

