## Supplemental Figures and Table for "Loss of NR2E3 disrupts rod photoreceptor cell maturation causing a fate switch late in human retinal development"

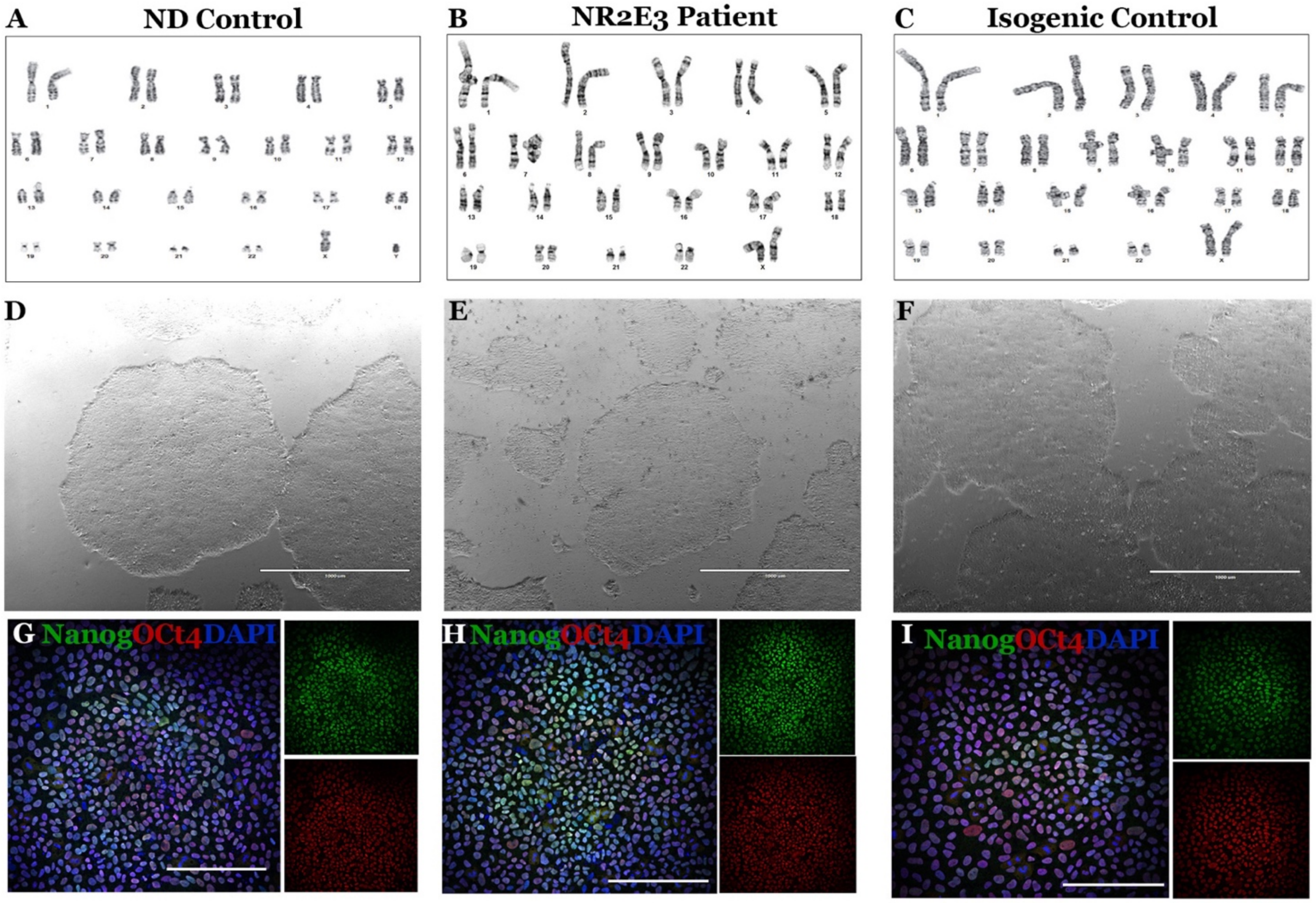


**Figure S1. Characterization of iPSC lines. A-C)** All iPSC lines displayed a normal karyotype by G-banding analysis. **D-F)** Patient and control lines displayed typical iPSC colony morphology throughout culture. **G-I)** Undifferentiated iPSC lines express markers of pluripotency including Nanog and OCT4.

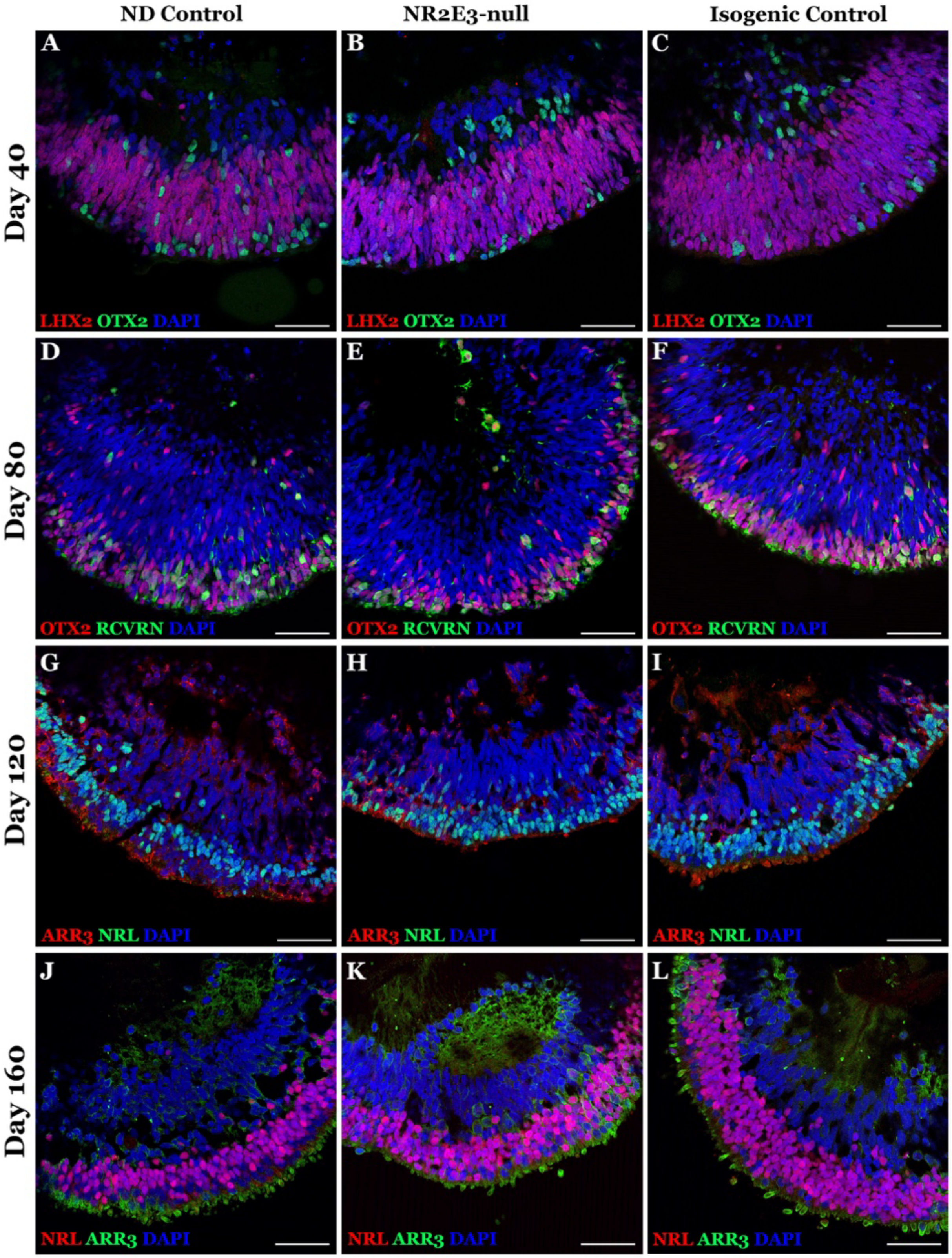


**Figure S2. Retinal organoid expression of normal differentiation marker genes. A-C)** Organoids from all three lines express *LHX2* and *OTX2*, markers of early neural commitment. **D-F)** By D80 of differentiation, the pan-photoreceptor marker *RCVRN* is expressed in organoid cells. **G-I)** At D120 of differentiation, the cone-specific arrestin *ARR3* and rod-specific factor *NRL* are expressed in the outer photoreceptor layer of organoids. **J-L)** By D160, organization of cone and rod photoreceptor layers is observed. Scalebars represent 50 µm.

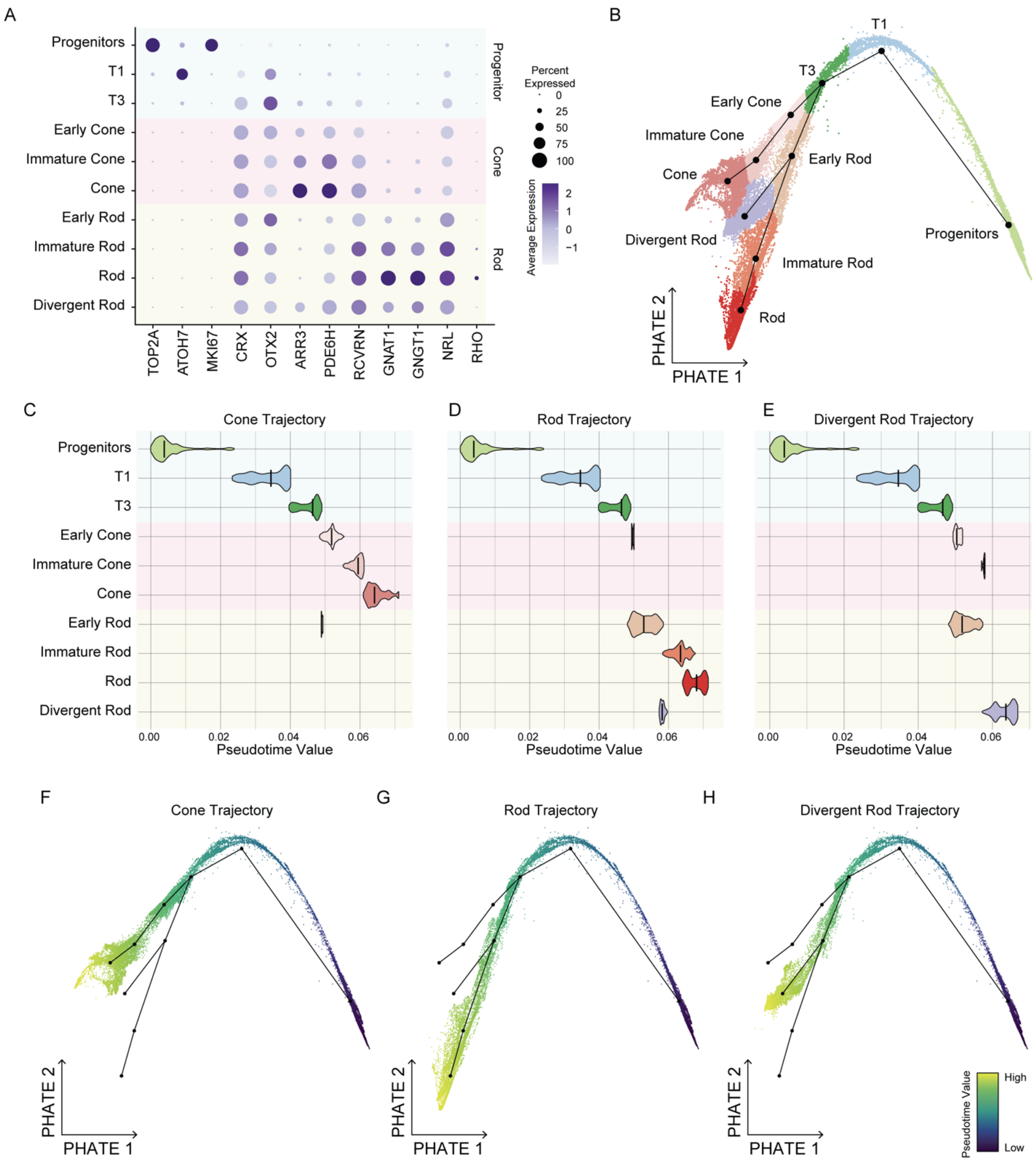


**Figure S3. PHATE reduction and trajectory analysis of photoreceptor lineage cells. A)** Clusters identified following PHATE reduction express appropriate progenitor-, cone-, and rod-specific genes. **B)** Annotated cells are shown based on PHATE dimensionality reduction. Branching lineages connecting clusters as identified by Slingshot are shown. Two branchpoints are identified giving rise to cones and rods and divergent rods respectively. **C-E**) Pseudotime values were generated for each cell using the three lineages identified in **B**. A violin plot is shown for each trajectory, showing the median pseudotime values (bar) for cells annotated for each cluster. Pseudotime values follow annotated clusters in terms of cell type maturity. Progenitor, T1, T3, and Early Rod clusters contain cells with comparable pseudotime values between the Rod and Divergent Rod trajectories (**D, E**). **F-H**) Trajectories based on the lineages in **B** are shown. Cells are colored based on pseudotime value along each trajectory with purple indicating a lower pseudotime value and yellow indicating a higher pseudotime value (arbitrary units).

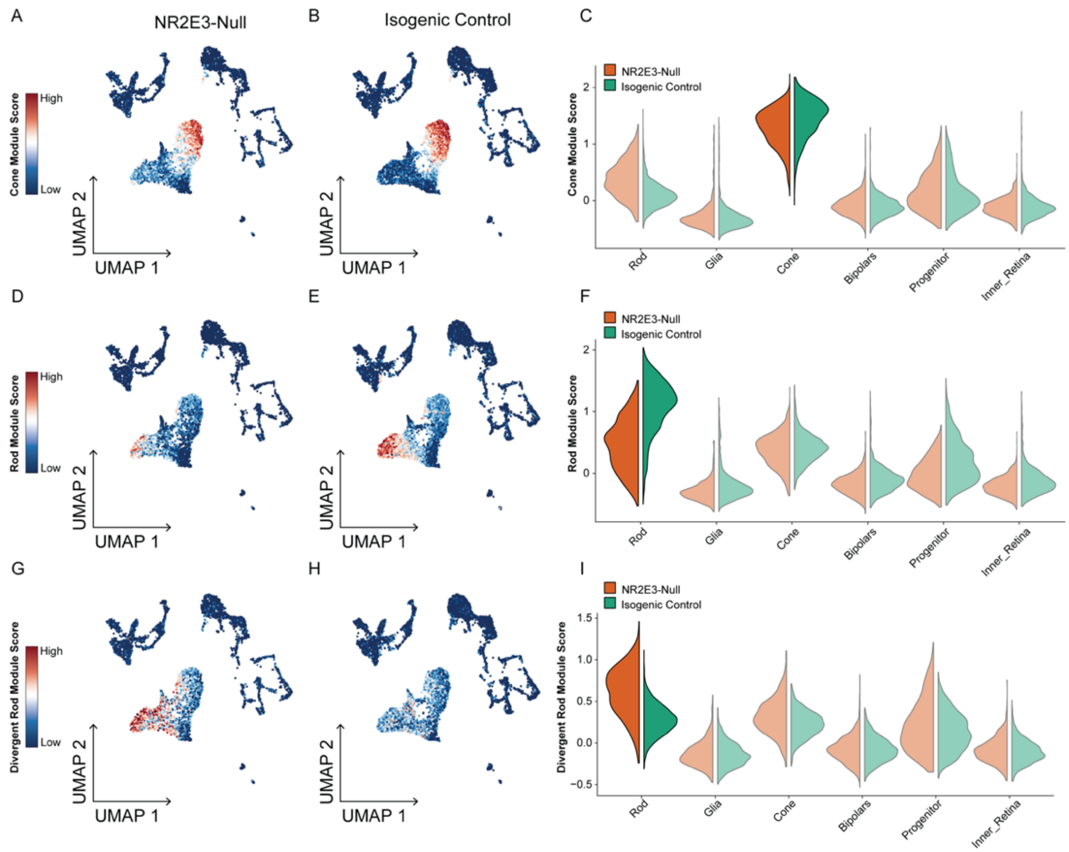


J

| Divergent Rod Module | Rod Module | Cone Module |
|---|---|---|
| VTN | GNAT1 | PDE6H |
| NR2E3 | GNGT1 | ARR3 |
| NSMCE2 | PDE6G | GUCA1B |
| SYNDIG1 | SAG | GUCA1C |
| AC112206.2 | CABP5 | AC112206.2 |
| AC063980.1 | RP1 | MYL4 |
| ISOC1 | NRL | SLC24A2 |
| NEUROG1 | ROM1 | CC2D2A |
| AC106798.1 | CNGB1 | CNGB3 |
| AC055733.2 | GNB1 | ASIC2 |
| SERPINF1 | PPEF2 | AC023905.1 |
| GNGT1 | GPR160 | KCNB2 |
| ROM1 | AIPL1 | RHOBTB1 |
| RCVRN | PDE6A | DPYSL3 |
| LINC00592 | DMD | MPP4 |
| AL391650.1 | YBX3 | PEX5L |
| HLA-C | CLUL1 | CCSER1 |
| FAM155A | MARCH1 | HCN1 |
| FAM174B | PPP2R2B | CAMK1D |
| TDRD7 | AP000462.1 | LINC02343 |
| AC104938.1 | PCDH15 | LRRC39 |
| KCNG4 | FAM161A | PRR16 |
| GUCA1B | ANK3 | NXNL1 |
| PAG1 | USH2A | RTBDN |
| WWC1 | PRPH2 | PPP4R4 |

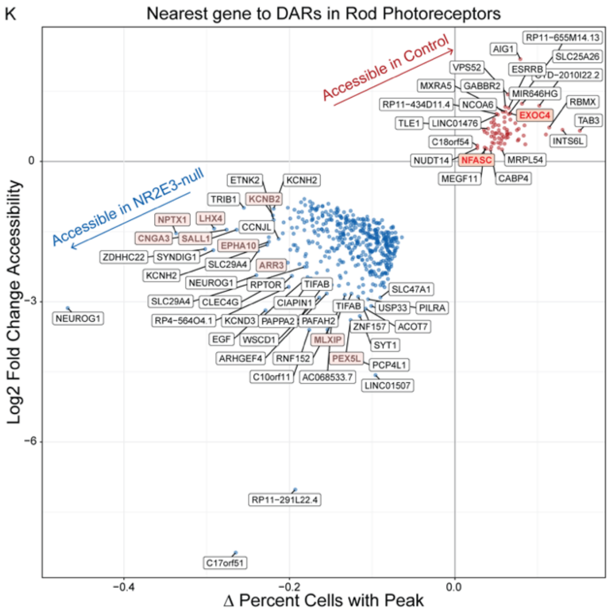


**Figure S4. Identification of divergent rods in multimodal sequencing data. A-B)** Cells assayed by multimodal sequencing are colored based on enrichment score for the cone gene module, with red indicating higher score. **C)** Cone gene module scores are plotted by cell type split by iPSC line of origin. Both NR2E3-null and Isogenic Control lines generate cones with high cone module score. **D-F)** As in **A-C**, cells are shown with a rod gene module score. There is notable enrichment of the rod module score in Isogenic Control rods versus NR2E3-null rods. **G-I**) The rod cluster of the NR2E3-null line shows the highest gene module score for the Divergent Rod module. **J)** Genes used to construct gene modules used in A-I. **K)** Differentially accessible regions between NR2E3-null and isogenic control rod photoreceptors (i.e., those from Figure 3D) are shown labeled with the symbol of the nearest gene. Within regions preferentially accessible in the NR2E3-null cells (blue dots), cone-specific genes are highlighted in pink. In regions preferentially accessible in control cells (red dots), rod-specific genes are highlighted in red.

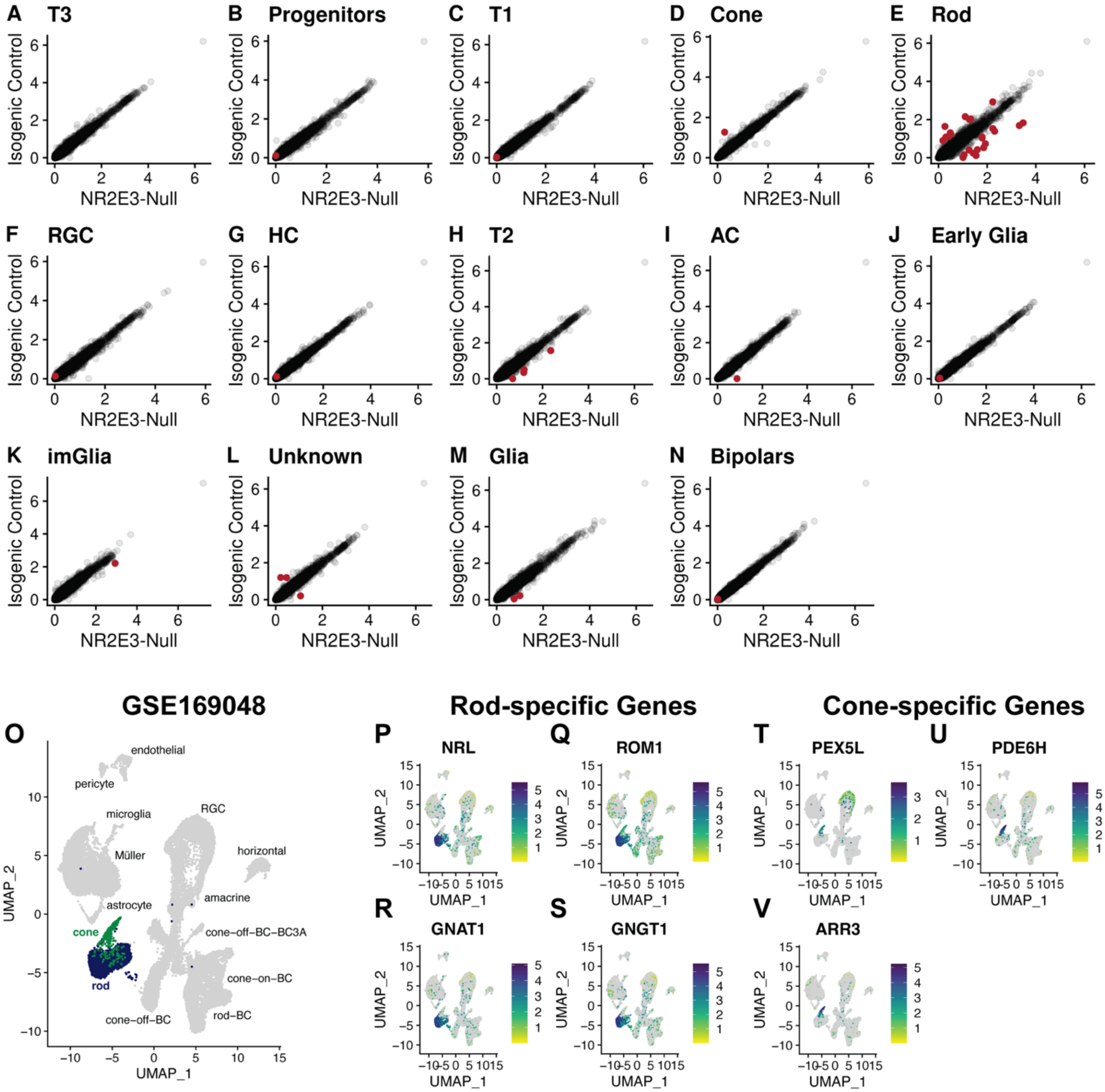


**Figure S5. Gene expression in retinal organoid and human donor retinal cells. A-N)** Dysregulation of gene expression in the NR2E3-null is restricted to rod photoreceptors. For each cell type, average expression level of every gene is shown on a log1p scale between the NR2E3-null and Isogenic Control lines. Significantly differentially expressed genes (i.e. Log2(FC) > 1 and delta % cells expressing > 10%) are shown in red. **O)** UMAP of single cells from human donor neural retinal cell types assayed by scRNASeq. Data was accessed from GSE169048. Rod and cone photoreceptors are indicated in blue and green, respectively. **P-S)** Expression of *NRL, ROM1, GNAT1,* and *GNGT1* is restricted to human rod photoreceptors. **T-V)** Expression of *PEX5L, PDE6H,* and *ARR3* is restricted to human cone photoreceptors.

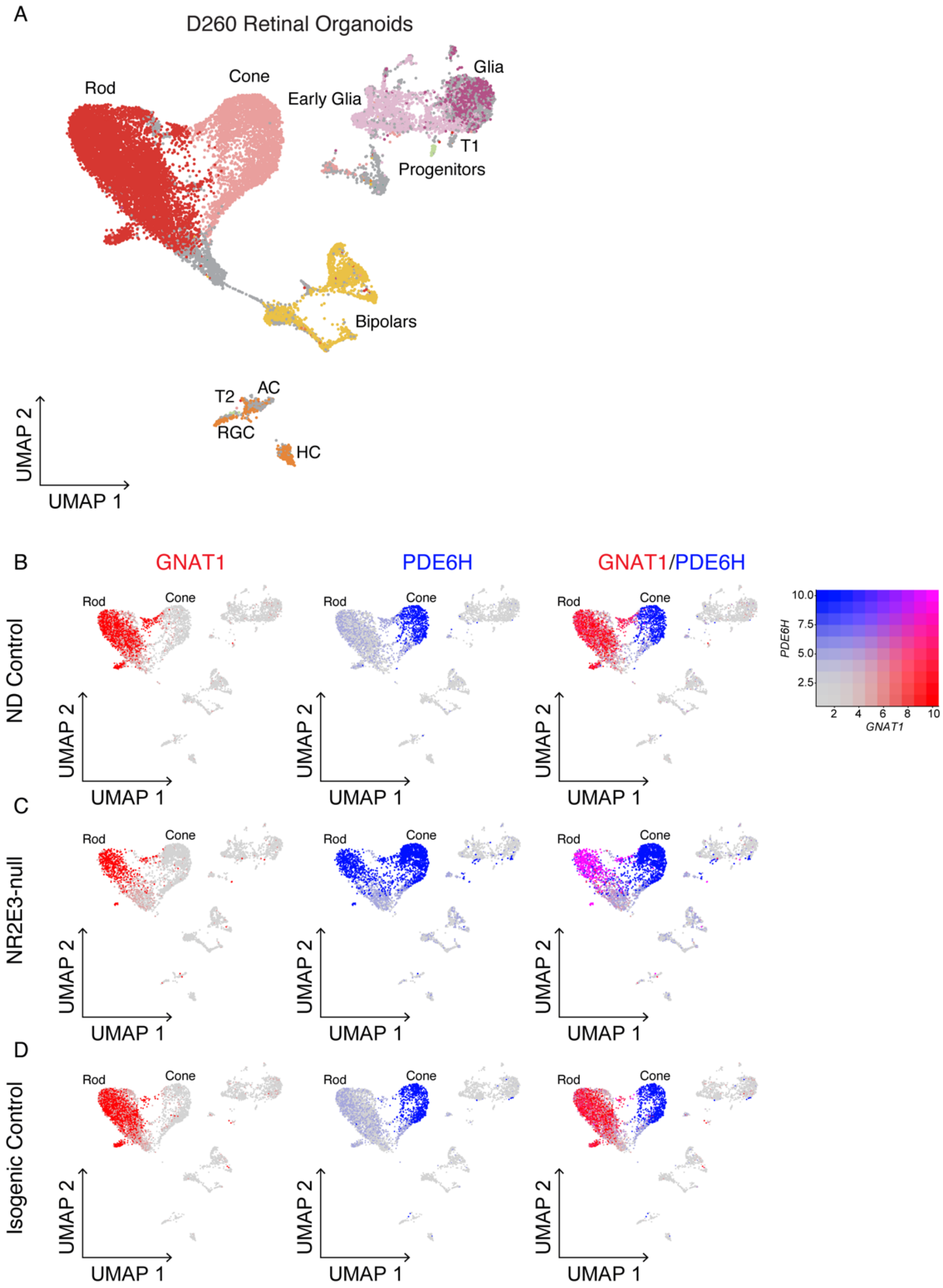


**Figure S6. scRNAseq in D260 retinal organoids shows persistence of divergent rods. A)** Cells from ND-control, NR2E3-null, and Isogenic Control lines collected at D260 of differentiation. Cells are projected in 2D space based on UMAP reduction derived from gene expression data. Late-stage organoids contain primarily photoreceptor, Müller glia, and bipolar cells. **B)** GNAT1, and PDE6H expression co-segregated into rod and cone photoreceptors in the ND control line. **C)** NR2E3-null organoids contain a population of photoreceptors that co-express GNAT1 and PDE6H (shown in pink, right panel). These represent the Divergent Rods described in the D40-D160 dataset. **D)** Monoallelic correction of NR2E3 restores normal segregation of GNAT1 and PDE6H expression in rod and cone photoreceptors.

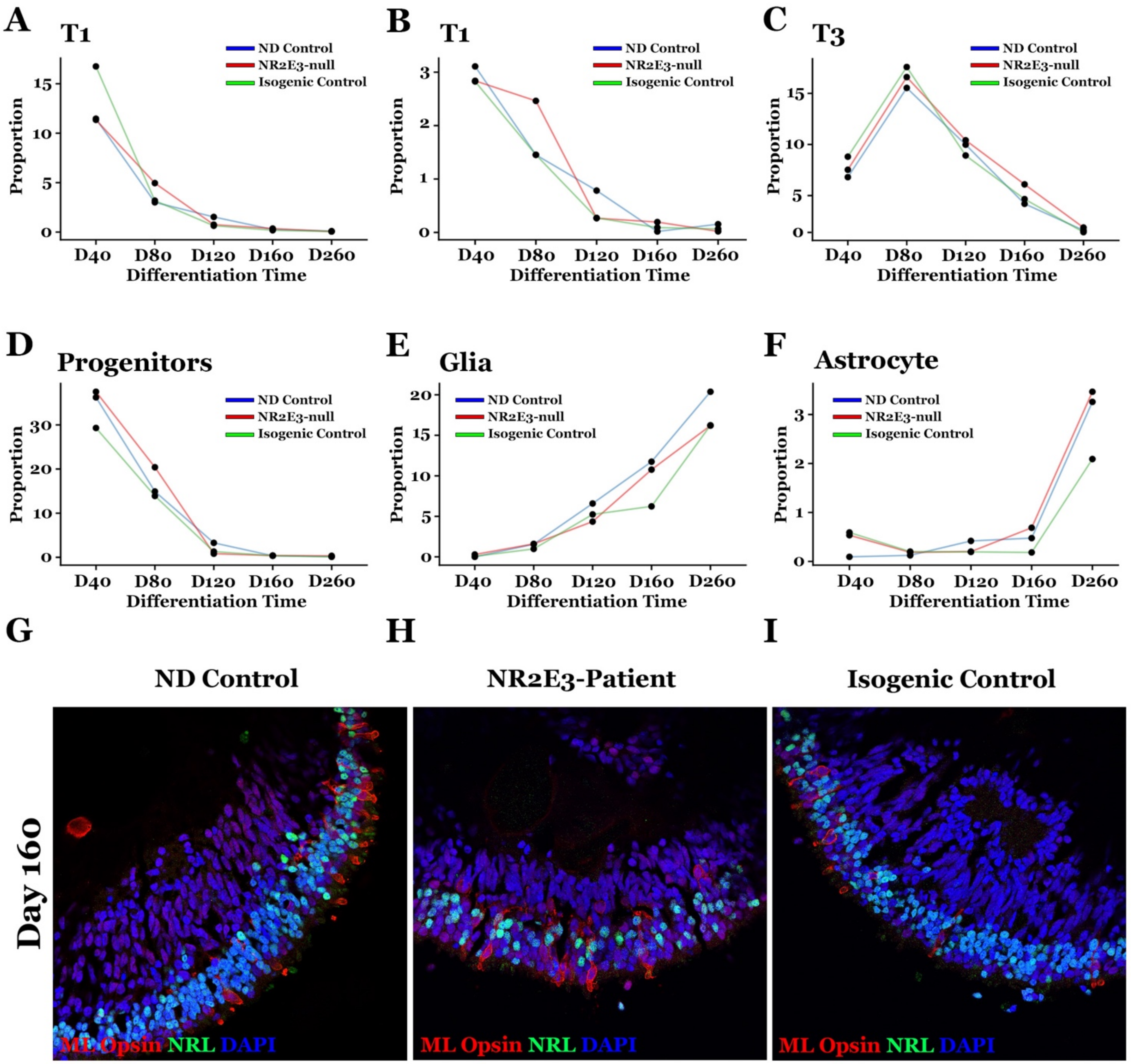


**Figure S7. Organoid composition across D40-D260 of differentiation. A-F**) The proportion of each cell type per timepoint within sampled organoids is shown. Between D160 and D260, no major differences are observed between lines. **G-I)** D160 organoids stained for the rod transcription factor NRL and the cone M/L-opsin. NR2E3-null (**H**) organoids do not contain a markedly greater number of M/L-opsin-positive cells than control organoids.

| **Table S1. Antibodies used.** | | | |
|---|---|---|---|
| ARR3 | Rabbit | LifeSpan Bio | LS-C368677 |
| CHX10 | Sheep | Exalpha | X1179P |
| Cleaved-PARP | Rabbit | Cell Signaling | 5625 |
| GNAT1 | Rabbit | Thermo | PA5-28336 |
| Ki67 | Rabbit | Abcam | AB15580 |
| LHX2 | Rabbit | Abcam | AB184337 |
| ML opsin | Rabbit | Millipore | AB5405 |
| Nanog | Goat | R&D Systems | AF1997 |
| NR2E3 | Mouse | R&D | PP-H7223-00 |
| NRL | Goat | R&D Systems | AF2945 |
| OCT4 | Rabbit | Stemgent | 09-0023 |
| OTX2 | Goat | R&D Systems | AF1979 |
| PDE6H | Mouse | Santa Cruz | SC-166350 |
| Recoverin | Rabbit | Millipore | AB5585 |
| Rhodopsin | Mouse | Millipore | MAB5316 |
| S opsin | Rabbit | Millipore | AB5407 |
| SNCG | Mouse | Abnova | H00006623-MO1 |
